# Genetic effects of plant–plant interactions depend on neighbor identity and environmental context

**DOI:** 10.64898/2026.07.28.741233

**Authors:** Noa Vazeux-Blumental, Carine Palaffre, Creola Brezeanu, Petre M. Brezeanu, Bernard Lagardère, Cyril Bauland, James Burridge, Pablo Affortit, Alexandre Grondin, Marzia Rossato, Andrea Benazzo, Tristan Mary-Huard, Laurence Moreau, Filippo Servalli, Roberto Papa, Elisa Bellucci, Elena Bitocchi, Laurent Laplaze, Domenica Manicacci, Maud I. Tenaillon

## Abstract

Plant–plant interactions shape community productivity and resilience, yet how genetic variation influences these interactions remains poorly understood. We conducted multi-environment field phenotyping of 200 common bean (*Phaseolus vulgaris*) lines grown in association with three maize (*Zea mays* ssp. *mays*) landraces to characterize the determinants of interspecific interactions. Our results reveal that maize landraces constitute distinct biotic environments, and interactions were strongly asymmetric, with maize acting as the dominant competitor. Bean responses to the identity of the maize partner involved phenology, vegetative development, productivity, and root architecture, and varied across field environments, with no evidence of bean local adaptation. We used genome-wide association to link variation across the bean genome to variation in bean and maize traits, thereby capturing both direct and indirect genetic effects, and tested for maize × bean interactions. We identified six genomic regions with indirect effects: three with stable effects across maize partners and three whose effects varied with maize partner identity, including candidate genes involved in neighbor perception and facilitative interactions. Our findings demonstrate that the phenotypic effects of genetic variation extend beyond individual plants, with competitive tolerance modulated by the genetic identity of neighboring plants and the environment.

**Highlight:** Neighbor identity shapes plant development and competitive tolerance through indirect genetic effects that are either stable or partner-dependent, revealing genomic regions potentially involved in neighbor perception and facilitative interactions.

## 1. Introduction

The positive impact of biodiversity on the functioning of plant communities is widely recognized and observed across temporal (Guerrero-Ramírez *et al.,* 2017) and spatial (Hautier *et al.,* 2018) scales. Functional complementarity among interacting species stands out as a key mechanism underlying these effects (Loreau and Hector 2001). From the perspective of a focal plant, the genotype and phenotype of its neighbors hence constitute a central biotic component of its environment. Plant–plant interactions range from competition to complementarity and facilitation and can operate both above and below ground (Duchene *et al*., 2017). Above ground, differences in architecture and phenology alter access to light and can modify radiation-use efficiency (Liu *et al*., 2018). Below ground, spatial segregation of root systems can modify nutrient acquisition (Postma and Lynch, 2012; Zhang *et al*., 2014), while rhizosphere processes and mycorrhizal networks can alter phosphorus and nitrogen availability (Li *et al*., 2007; Gutiérrez-Núñez and Gavito, 2024). Together, these processes show that neighboring plants are not passive components of the environment: they can modify the resources and conditions experienced by one another, with consequences for growth, architecture, and reproduction.

While the phenotypic consequences of plant–plant interactions are well documented, much less is known about how genetic variation within interacting species contributes to these responses. Direct genetic effects describe the influence of an individual’s genotype on its own phenotype, whereas indirect genetic effects describe the influence of neighboring genotypes on the focal individual’s phenotype (Moore *et al*., 1997; Wolf *et al*., 1998). An open question is whether indirect genetic effects are consistent across neighboring genotypes or depend on partner identity, potentially generating genotype × genotype (G × G) interactions that may themselves vary across abiotic conditions (G × G × E). This complexity has recently been dissected through quantitative approaches (Sato and Wuest, 2025), including co-genome wide association (co-GWA), which identified interacting loci affecting shade avoidance in wheat variety mixtures (Mathieu *et al*., 2024, Preprint). Comparing indirect genetic effects across contrasting partner genotypes can also help distinguishing loci with stable effects from those whose effects depend on partner identity.

Cereal–legume mixtures provide a particularly relevant system in which to investigate plant-plant interactions, because the component species differ strongly in architecture, resource acquisition, and competitive ability. In maize–bean intercropping, for example, complementarity and facilitation have been reported for light, soil nitrogen, phosphorus, and root foraging (Postma and Lynch, 2012; Zhang *et al*., 2014; Gutiérrez-Núñez and Gavito, 2024), while competition can strongly constrain bean development (Baudoin *et al*., 1997; Vazeux-Blumental *et al*., 2025). This contrast in ecological strategies makes maize–bean mixtures particularly suited to examine how variation in neighbor identity affects the phenotype and genetic architecture of a competition-sensitive focal species, from above-ground development to root architecture.

Maize (*Zea mays* L. subsp. mays) and common bean (*Phaseolus vulgaris* L.; hereafter bean) also share a long history of association. Both were domesticated in Mexico (Chacón *et al*., 2005; Piperno *et al*., 2009), where their cultivation together with squash gave rise to the milpa system (Zizumbo-Villarreal *et al*., 2012), and both were introduced into Europe in the 15^th^ century (Brandenburg *et al*., 2017; Bellucci *et al*., 2023). In some European regions, such as Transylvania, maize–bean intercropping has persisted as a traditional practice (Vazeux-Blumental *et al*., 2024). This history raises the question of whether long-term association influences partner compatibility, as reflected in the relative productivity of the two component crops in mixtures.

We used 200 European climbing bean lines, including lines with a history of traditional co-cultivation with maize, and three maize landraces. We evaluated bean response across four field experiments spanning two years and two countries, combining above-ground developmental and productivity traits with below-ground root phenotyping. By integrating multi-environment field phenotyping with GWA, we asked whether maize partner identity alters bean phenotypes, whether interspecific effects are asymmetric and consistent across environments, and whether association history predict partner compatibility. Finally, we tested whether indirect genetic effects depend on partner identity.

## 2. Materials and methods

### 2.1. Panel design and initial seed production

#### 2.1.1. Initial bean panel genotyping

We relied on a panel of 637 climbing common bean (thereafter bean) lines previously genotyped (Bellucci *et al*., 2023). Those lines were derived from European landraces through one to four generations of selfing (Single Seed Descent, SSD, Table S1). Because bean is predominantly self-pollinating, this was sufficient to ensure genetic homogeneity, with offspring being homozygous at most loci. In addition, we assembled a new collection of 57 lines consisting of on-farm collected landraces, including 37 Romanian, 6 French, and 14 Italian accessions (Table S1). Those landraces have historically been intercropped with maize and thereafter referred to as co-cultivated lines. Note that these lines displayed greater genetic heterogeneity than the SSD lines.

DNA for the 57 new lines was extracted from young leaflets of single plants using the DNeasy 96 Plant Kit (Qiagen GmbH, Hilden, Germany). As for the initial 637 lines, we used Genotyping-by-Sequencing (GBS; Elshire *et al*., 2011) with a two-enzyme system (TaqαI and MseI) optimized according to Schröder *et al*. (2016). After PCR amplification, samples were purified with AMPure XP Beads, and libraries were sequenced on an Illumina platform at the Department of Biotechnology, University of Verona (Italy).

All data from the 694 lines thereafter referred to as the “initial bean panel” were analyzed jointly. Low-quality reads were filtered out and trimmed using Cutadapt v3.2 (Martin, 2011), and read quality was assessed with FastQC (http://www.bioinformatics.babraham.ac.uk/projects/fastqc/). Cleaned reads were mapped to the *Phaseolus vulgaris* v2.0 reference genome (Schmutz *et al*., 2014) using BWA-MEM v0.7.15 (Li, 2013, Preprint). Aligned reads were sorted and filtered for optical duplicates with Picard v2.4.1 (http://broadinstitute.github.io/picard). Variant calling was performed with GATK v4.1.9.0 (McKenna *et al*., 2010) using the HaplotypeCaller for each accession and GenotypeGVCFs for joint genotyping. SNPs were extracted with SelectVariants and filtered using VariantFiltration with the following hard thresholds: QUAL < 60 || QD < 2.0 || MQ < 40.0 || FS > 60.0 || SOR > 3.0 || MQRankSum < -20.0 || ReadPosRankSum < -8.0. INDELs and heterozygous genotypes were replaced by missing data.

#### 2.1.2. Genetic structure of the initial bean panel

We assessed the genetic structure of the initial bean panel and used it to select lines included in our intercropping panel. Prior to analysis, GBS data were filtered using VCFtools, Plink, and Tassel (*Phaseolus vulgaris* v2.1) scaffolds, non-biallelic and heterozygous SNPs were removed; SNPs with depth <3×, MAF <5%, >30% missing data, or <10 kb apart were discarded; individuals with >50% missing data were excluded. We performed a supervised ADMIXTURE analysis (Alexander *et al*., 2009) using 20 Mesoamerican (MA) and 20 Andean (A) reference lines of American origin, previously shown to be non-admixed (Bellucci *et al*., 2023, Table S2). From the admixture results, we retained 317 lines with a MA ancestry >80%, as well as seven co-cultivated Italian lines from the A gene pool (Table S2) as candidates for the intercropping experiments.

#### 2.1.3. Seed production and intercropping panel

Bean seed multiplication was performed to ensure that the GWA field trials rely on a single seed lot per line. To do so, the 317 lines were grown in 2021 in an open field either at the INRAE station of Saint-Martin-de-Hinx (SMH) or in greenhouses at Maugio (INRAE, Southern France) or Orsay (Université Paris-Saclay) for seed production. From these, a panel of 200 bean lines was selected for intercropping based on seed availability, including 179 SSD lines and 21 co-cultivated lines (8 Romanian MA, 6 French MA, 7 Italian A).

Three genetically heterogeneous maize landraces were selected (Table S1): Grand Roux Basque (France), Spinato di Gandino (hereafter Gandino, Italy), and Văleni (Romania). All are used traditionally for human consumption (e.g., polenta, flour for bread and cakes). The Văleni seeds were collected on-farm (with intercropped beans), and registered at the Institute of Biological Research Cluj-Napoca (doi:10.18730/10CJN5). Maize seeds were produced through controlled crosses at SMH for Văleni and at Agricola Clemente Savoldelli for Gandino (Italian national variety registration No. 16342). The Grand Roux Basque seed lots came from the CRB Gamèt located in Montpellier as a mix of two seed lots (FRA0410610 and FRA0410641), both predominantly assigned to the Pyrenean– Galician genetic group (Arca *et al*., 2023).

### 2.2. Field assays and phenotyping of the intercropping panel

#### 2.2.1. Experimental design

The assays took place at Saint-Martin-de-Hinx (hereafter SMH) in France (3°34’19.2"N; 1°18’0"W at 30 m altitude) and at the experimental station of Bacău in Romania (46°35’6.74"N; 26°57’0.31"E at 165 m altitude). They were replicated in 2022 and 2023. The soil at the French site consists of clay-loamy sand (luvisol), while the soil at the Romanian site is a well-developed, loamy-sandy textured polished cambic chernozem (fluvisol). Each field included two blocks, each divided into three sub-blocks corresponding to the three maize landraces (Fig. 1). Within each sub-block, the 200 bean lines were sown directly alongside the corresponding maize, in rows of 30 plants (15 maize and 15 beans), resulting in a total of 1,200 plots per country and year (200 bean lines × 3 maize landraces × 2 blocks).

**Figure 1.**
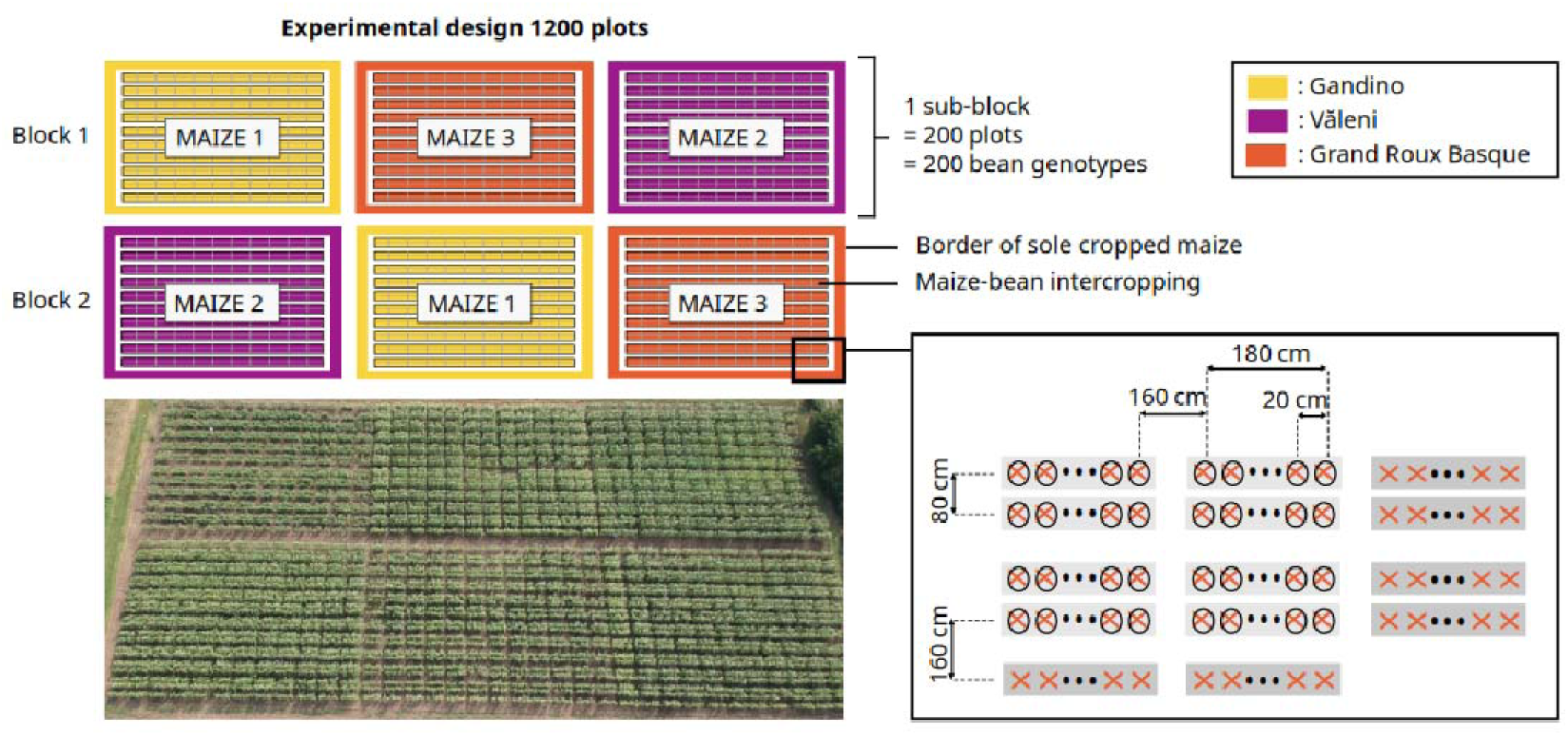
Experimental design comprising 1,200 maize–bean intercropped plots, where 200 bean lines were associated with three maize landraces. The overall design (top left) includes two blocks, each divided into three sub-blocks corresponding to the maize landraces (shown in distinct colors), bordered by sole-cropped maize. Within each sub-block, 200 bean lines were randomly assigned to plots containing 15 maize and 15 bean plants (bottom right). The bottom-right diagram illustrates eight intercropped plots with MAIZE3 and various bean lines (light grey) and seven sole-cropped maize plots (border plots in dark grey) (maize = crosses; beans = circles). The experiment was conducted in France and Romania in 2022 and 2023. The bottom-left photograph shows the Romanian field in 2022. ALT TEXT: Schematic of the randomized two-block maize-bean field design used in France and Romania, with three maize-landrace sub-blocks per block, 200 bean lines randomized within each sub-block, and a photograph of the 2022 Romanian field.

Sub-blocks were bordered by maize sole-crop plots of the same maize landrace, and both subblocks and rows were randomized within countries and years. Plant spacing was 20 cm within rows, measured between each maize–bean plant pair, and 80–160 cm between rows, reflecting typical maize-bean intercropping densities used in France (Vazeux-Blumental *et al*., 2025). Maize and bean were sown at 6 cm and 2 cm depth, respectively, on the same day between May 8^th^ and June 7^th^ in France and in Romania in 2022, while in Romania in 2023, beans were sown three weeks after maize (May 16th).

The four field trials were carried out in low-input settings, and bean seeds were inoculated with a symbiotic rhizobia mix (LEGUMEFIX *Phaseolus* Inoculant, UK). Fertilization was applied as N < 58.5, P < 104.5, K < 108. Three irrigation events occurred in Romania in 2022 (92.86 mm total), and no irrigation was applied in other sites. Grand Roux Basque was not sown in Romania in 2023, due to lodging in 2022. Also, because of a lack of seeds, one bean line was replaced in 2023, and border rows of maize Văleni were replaced by a commercial variety in Romania in 2022.

#### 2.2.2. Above-ground plant phenotyping

Across all fields, 12 to 15 morphological and agronomic traits were measured on bean plants, and 7 to 8 traits on maize plants (Table S3). Thermal times were calculated from sowing to emergence, to flowering, and to pod formation, and were expressed in degree days in Celsius (°Cd) as the cumulative difference between the daily mean temperature and the base temperature for vegetative growth (10°C for beans and 6°C for maize) according to Bonhomme (2000).

Two additional traits were computed from maize and bean yields to quantify overall mixture performance. Let □denote the set of all observed maize–bean combinations (*k,l*) within an experimental field, where *k* indexes maize landraces and *l* indexes bean lines. For a field plot containing maize landrace *i* and bean line *j*, *M_ij_* and *B_ij_* denote the total maize kernel weight and total bean seed weight, respectively. The parameters μ*_M_* and σ*_M_* denote the mean and standard deviation of maize yields across all plots within the experimental field, while μ*_B_* and σ*_B_* denote the corresponding mean and standard deviation of bean yields.

We first standardized maize and bean yields as:

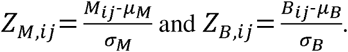

The cumulated yield index was then calculated as:

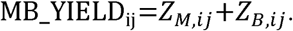

To define the balance-penalized performance index, we first calculated:

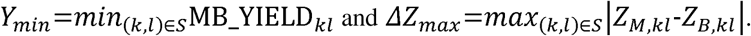

The balance-penalized index was then defined as:

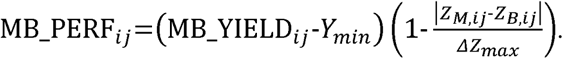

*MB_YIELD_ij_* measures the combined standardized production of maize landrace *i* and bean line *j*, whereas *MB_PERF_ij_*additionally penalizes imbalances between the standardized yields of the two species. Thus, combinations with similar maize and bean standardized yields receive a lower penalty than combinations dominated by one species. Finally, for each bean line *j*, mean *MB_PERF* was calculated across the three maize landraces within each experimental field.Because bean was the focal species, maize traits were measured on subsets of plots whose number varied according to the country and year. In France, all maize traits were assessed across the 1,200 plots in 2022 and 2023, except for three traits (thermal time to 50% anthesis, total plant height, flag leaf width) measured on 258 plots in 2023 (Table S3). In Romania, in 2022 plant height and thermal time to 50% anthesis were measured on all 1,200 plots, while the other traits were assessed on 228 plots; in 2023, all eight traits were measured on 172 plots (Table S3).

#### 2.2.3. Below-ground phenotyping

Below-ground phenotyping was conducted in France in 2022 and 2023 at the flowering stage of beans (40–45 days after sowing), representing the peak for plant vigor. We phenotyped roots of beans in all 600 plots (200 bean lines × 3 maize) in a complete block, while in the second block, we assessed a subset of ∼40 bean genotypes (plots) per sub-block (120–150 additional plots). Note that these 40 plots were chosen to match those used for maize above-ground phenotyping, enabling the computation of correlations between shoot and root traits. Only three of the 15 bean plants per plot were phenotyped due to the destructive nature of root measurements. These plants were selected at the plot edges so as not to affect subsequent aerial measurements. Maize roots were measured from sole-cropped border plants in 2022 and from both border plots and intercropped plants in 2023; note that border plots had half the plant density of intercropped plots.

In both species, roots were gently excavated with a shovel, rinsed thoroughly, and carefully separated by hand during washing to untangle intertwined roots. Phenotyping combined manual and image-based approaches: nine traits were measured on beans (seven related to root architecture, one to nodulation, one to disease incidence), and seven traits were measured on maize (Table S3). Manual bean phenotyping was performed on a single plant per plot, whereas image-based measurements included all three plants.

### 2.3. Statistical analysis

Statistical analyses were conducted using *R v4.4.1* (R Core Team, 2024). Statistical models were fitted using the *R* packages *Sommer* (Covarrubias-Pazaran, 2016) and *lme4* (Bates *et al*., 2015). To assess the significance of the effects, we used likelihood ratio (LR) tests for the linear mixed models, and F-tests from ANOVA fixed-effects linear models.

In the models described below, μ stands for the model intercept and ε for the residual error term, assumed to follow normal distribution *N(0,* σ*²E)*, *Rk* for the fixed effect of block *k*, *Mi* for the fixed effect of maize *i*, *Cl* for the fixed effect of country *l*, *Ym* for the fixed effect of year *m*, *Gj* for the random effect of bean genotype *j*, assumed to follow *N(0,* σ*²G)*. Interaction terms are defined in the same way as the corresponding main effects. Interactions modeled as random effects are considered to follow normal distributions with specific variance for each maize (in *G* × *M*), each country (in *G* × *C*), and each year (in *G* × *Y*).

#### 2.3.1. Broad-sense heritability in each field

We estimated broad-sense heritability (*H²*) for each bean and maize trait (Visscher *et al*., 2008) using a linear mixed model within each country, year, and maize environment for above-ground traits, and within each year and maize environment in France for below-ground traits:

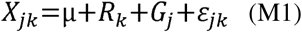

where *Xjk* is the phenotypic value (measured on bean or on maize) for a plot sown with the bean genotype *j* in block *k*.

Heritability was calculated as:

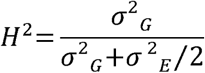

with σ*^2^G* the genotypic variance associated with the random effect *Gj*, and σ*^2^E* the residual variance; the residual variance is divided by 2 because each genotype was evaluated in two blocks.

#### 2.3.2. Spatial corrections within field

Corrections were performed for the block effect and for maize × block interaction effects on quantitative variables by fitting the same linear mixed model (Laird and Ware, 1982) for both maize and bean datasets and for the four experimental fields (2 countries and 2 years) separately:

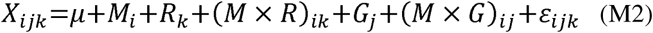

where *Xijk* is the maize or bean phenotypic value for a plot sown with maize *i* associated with bean genotype *j* in block *k*.

Maize and bean data were corrected for both *R* and (*M* × *R*) effects by subtracting these effects to the phenotypic values, yielding two corrected values per maize-bean combination (one for each block) per country and year. Due to limited root data, the (*M* × *G*) interaction was not tested for below-ground traits. For root traits measured manually, an additional manipulator fixed effect was included in the model, and data were corrected accordingly.

#### 2.3.3. Testing country and year effects

To test for year and country effects on above-ground traits, we fitted two global models on corrected maize and bean data (c.f. model M2).

For maize:

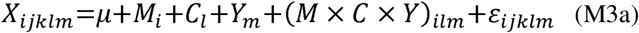

where *Xijklm* is the corrected phenotypic value for maize *i*, bean *j*, block *k*, country *l*, and year *m*.

For bean:

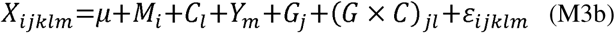

where *Xijklm* is the corrected bean phenotypic value for a plot sown with maize *i*, bean *j*, block *k*, country *l*, and year *m*.

#### 2.3.4. Correction for year effect in both countries

To correct for year effect on above- and below-ground traits, two models were fitted per country on data already corrected for block effect and block × maize interaction (model M2).

For maize:

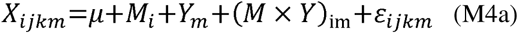

where *Xijkm* is the corrected maize phenotypic value for a plot sown with maize *i*, bean j, block k, and year *m*.

For bean:

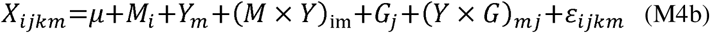

where *Xijkm* is the corrected bean phenotypic value for a plot sown with maize *i*, bean genotype *j*, block *k*, and year *m*. Note that (*Y* × *G*)*mj* was not included in the model for below-ground traits due to limited root data. We corrected further for year effect and year × maize interaction, and computed country-specific mean values, yielding a single value per maize-bean combination in each country.

#### 2.3.5. Bean response to maize biotic environment and interaction between species

We tested whether bean responses differed among the three maize landraces, considering them as biotic environments. First, we characterized differences among maize landraces using Tukey’s HSD tests and ANOVA from models M4a and M4b. Within-maize variability was assessed using variance estimates and Brown–Forsythe tests. Then we analyzed bean response to maize variability. We first illustrated the global effect of maize on bean phenotypes through a Principal Component Analysis (PCA). Bean response to maize was further evaluated in each country using model M4b and Tukey’s HSD applied to pairwise comparisons among the three maize landraces.

To characterize maize × bean interactions, pairwise Pearson’s correlations were computed between maize and bean traits, with significance assessed using a 5% false discovery rate (FDR; Benjamini and Hochberg, 1995). Except for phenology traits, negative and positive correlations were interpreted as interspecific competition and beneficial interactions, respectively. Finally, for each maize, we identified bean lines as top-performer partners if their yield was higher than the median yield across all beans and if the yield of the maize they were paired with was higher than the median yield for that maize associated with all beans.

### 2.4. Bean local adaptation

To investigate local adaptation in beans, we analyzed the relationship between bean phenotypes in each country and climatic distances between their collection site and the experimental field site. Nineteen bioclimatic variables (Table S4) were extracted for 183 of the 200 bean lines (Fig. 2) at 30-arc-sec resolution from the *CHELSA v2.1* database (Karger *et al*., 2017) for the 1982–2010 period. Climate variation across sites of collection was summarized by a PCA. For each of the four experimental fields, i.e., country × year, we computed Mahalanobis distances between the 183 collection sites and the experimental field site using 18 independent bioclimatic variables excluding bio7, which was a linear combination of other variables.

**Figure 2.**
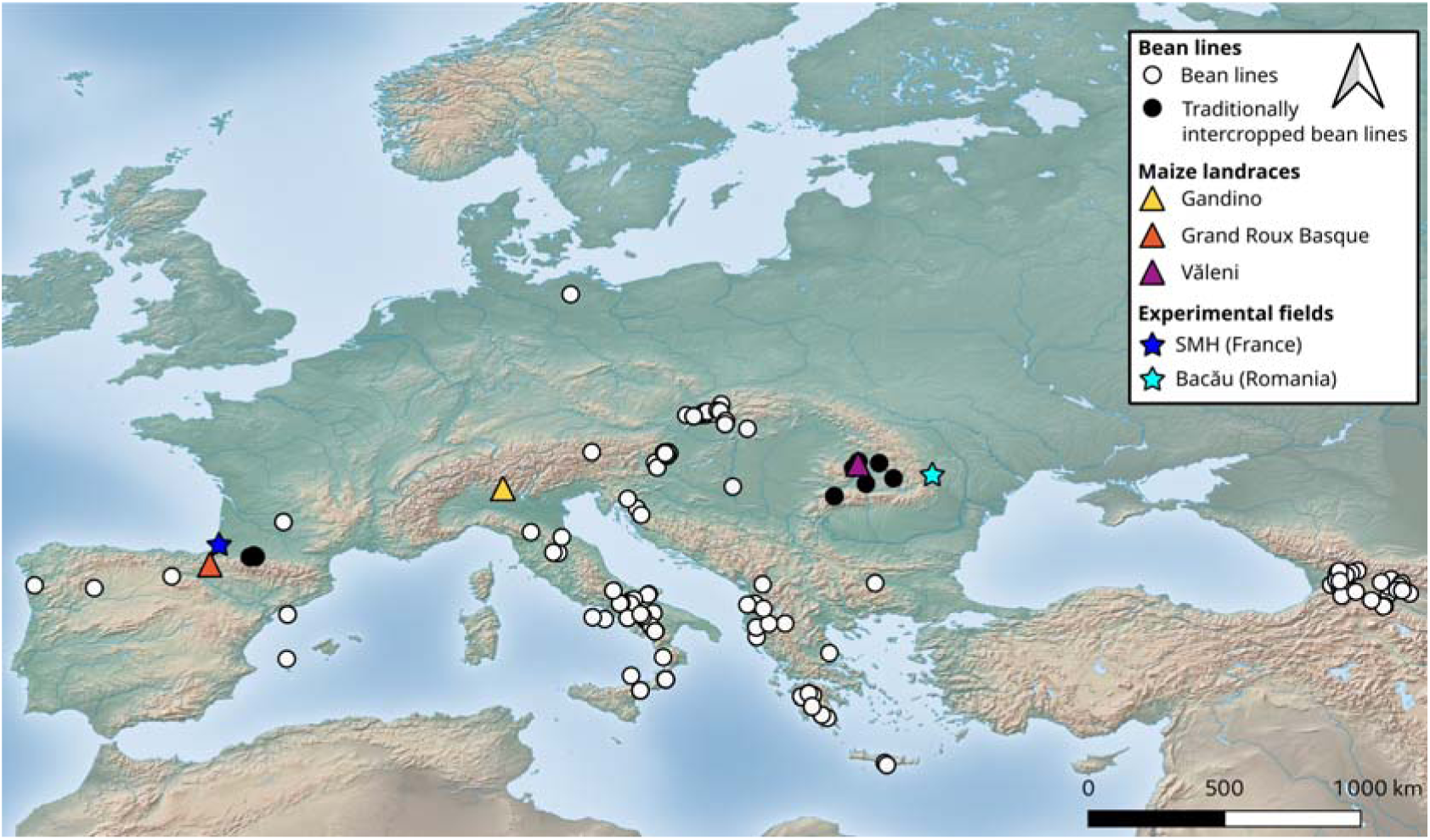
Geographical origin of bean and maize germplasm involved in the experiments, and location of field trials. Dots indicate the localization of origin of 183 European beans, all from Mesoamerican origin, including 21 lines traditionally used in intercropping with maize (black); the other lines are indicated in white. Triangles indicate the geographical origin of the three maize landraces used in the study. Stars indicate the localization of the two experimental sites (SMH in France and Bacău in Romania). ALT TEXT: Map of Europe showing the origins of the 183 Mesoamerican bean accessions including the subset traditionally intercropped with maize, the origins of the three maize landraces, and the French and Romanian field sites.

We then tested the correlation between bean traits and climatic distances using the 183 lines. Additionally, we regressed bean phenotypic distances on climatic, geographic, and genetic distances through multiple regression on distance matrices (MRM), using the R package *ecodist* with 1,000 permutations and Q-value < 0.05 (FDR calculated within each country and year).

### 2.5. Genome-wide association mapping on the bean genome

From the raw GBS data, SNPs with a sequencing depth below 3×, more than 30% missing data were discarded. SNPs with a minor allele frequency (MAF) < 5% were excluded from the direct genetic effect analysis. For the indirect genetic effect analysis, which relied on a smaller maize dataset, a MAF threshold of 25% was applied to ensure sufficient representation of the minor allele. INDELs and heterozygous genotypes were replaced by missing data. Genotypes were imputed using a reference panel of 6,007 *P. vulgaris* accessions with *Beagle v5* (Browning *et al*., 2018). Redundant SNPs were pruned for linkage disequilibrium (LD, r² > 0.99) within sliding windows of 50 SNPs, resulting in a final dataset of 11,142 SNPs for direct genetic effect and of 3,004 SNPs for indirect genetic effect.

Genome-wide associations were conducted using a mixed linear model (Yu *et al*., 2006) implemented in the *GAPIT* R package, applying a leave-one-chromosome-out approach. Analyses were performed separately for each maize, country, and year, resulting in a total of 5 to 11 GWA depending on phenotypic traits. Phenotypic data consisted of mean values per country corrected for the block effect using model M2. We conducted a joint analysis of the GWA outputs using the fixed-effect meta-analysis model implemented in the *metaGE* R package (De Walsche *et al*., 2025). This meta-GWA approach was employed to identify QTLs with stable effects across maize partners and country–year field environments. Multiple testing was handled through the local score (LS) procedure (Fariello *et al*., 2017) implemented in metaGE, using a P-value threshold of 2 (log10 scale) and a LS threshold fixed at 10 for direct genetic effects. No LS threshold was applied for indirect effects owing to the lower statistical power resulting from scarce maize phenotypic data. Additionally, the meta-GWA approach was applied to identify QTLs showing contrasting effects across biotic environments, identified using contrasts. The method was applied to bean and maize phenotypic traits to evaluate direct and indirect genetic effects, respectively. The fixed effect model was first used to identify QTLs with a common non-zero effect across environments by combining signed z-scores while accounting for correlations between environments. Within this model, the following contrasts were tested:

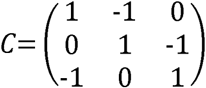

corresponding to Gandino versus Grand Roux Basque, Grand Roux Basque versus Văleni, and Gandino versus Văleni. Significant contrast tests indicated SNPs whose effects differed among maize landraces.

### 2.5. GO term enrichment and comparative annotation of candidate genes

We extracted the candidate genes under the QTL regions identified for each meta-GWA and performed gene ontology (GO) term enrichment using the *topGO* v2.56.0 R package (Alexa and Rahnenführer, 2024). Orthologous genes corresponding to the *P. vulgaris* candidate genes retrieved were identified using *OrthoFinder* v2.5.5 (Emms and Kelly, 2019). The analysis was performed using the primary-transcript protein sequences downloaded from Phytozome for *Arabidopsis thaliana* Araport11, *Cicer arietinum* v1.0, *Glycine max* Wm82.a2.v1, *Lens culinaris* v1, *Lupinus albus* v1, *Vigna unguiculata* v1.2, *Phaseolus vulgaris* v1.0, *Arachis hypogaea* Line8 v1.3, and *Pisum sativum* var. ZW6 v1.0. OrthoFinder output was subsequently parsed using a custom script to intersect the inferred orthologous relationships with the input list of *P. vulgaris* genes and retrieve the corresponding orthologs in the target species. We further conducted a comparative literature survey to identify *Phaseolus vulgaris* genes whose orthologs in other legume species or *Arabidopsis thaliana* have been reported or proposed to play a role in plant–plant interactions.

## 3. Results

### 3.1. Pervasive genotype × environment interactions

We analyzed bean and maize phenotypes across all experimental fields and maize landraces (two countries × two years × three maize landraces). Broad-sense heritability was first estimated separately within each environment from unadjusted data and then summarized across environments with mean and standard deviation (Table 1). For traits measured on maize, heritability ranged from 0.24 for total kernel mass per plant to 0.77 for main ear width, with high between-environment variation for most traits (up to a SD of 0.42 for plant height). Bean traits generally showed higher heritability, ranging from 0.41 for leaflet length to 0.96 for seed length.

**Table 1.** Broad-sense heritability mean and standard deviation (SD) for bean and maize traits.

| Species | Trait□ | Mean heritability (SD)□ |
|---|---|---|
| Bean | B_MEAN_LENGTH_LEAFLET | 0.41 (0.22) |
|  | B_MEAN_WIDTH_LEAFLET | 0.42 (0.26) |
|  | B_TOT_SEED_MASS_PLT | 0.45 (0.17) |
|  | B_FOLIAGE_DENSITY | 0.50 (0.19) |
|  | B_POD_PLACEMENT | 0.53 (0.27) |
|  | B_NBR_PODS_PLT | 0.54 (0.08) |
|  | B_TOT_NBR_SEED_PLT | 0.56 (0.10) |
|  | HEIGHT_B_OVER_M | 0.66 (0.09) |
|  | B_NBR_SEED_PER_POD | 0.66 (0.09) |
|  | B_FLOWERING_50_DD | 0.80 (0.14) |
|  | B_PODS_FORMATION_50_DD | 0.84 (0.07) |
|  | B_100_SEED_MASS | 0.91 (0.03) |
|  | B_SEED_WIDTH | 0.94 (0.02) |
|  | B_SEED_SURFACE | 0.96 (0.02) |
|  | B_SEED_LENGTH | 0.96 (0.02) |
| Maize | M_TOT_K_MASS_PER_PLT | 0.24 (0.30) |
|  | M_WIDTH_LEAF | 0.25 (0.04) |
|  | M_NBR_EARS_PER_PLT | 0.28 (0.18) |
|  | M_TOT_NBR_K_PER_PLT | 0.34 (0.37) |
|  | M_TKW | 0.50 (0.41) |
|  | M_LENGTH_MAIN_EAR | 0.60 (0.37) |
|  | M_HEIGHT_PLT | 0.61 (0.42) |
|  | M_FLOWERING_50_DD | 0.76 (0.36) |
|  | M_WIDTH_MAIN_EAR | 0.77 (0.08) |

□: Traits are described in Table S3
□: The mean and SD were computed across 6 to 11 environments (2 years x 2 countries x 3 maize landraces) for bean traits and across 2 to 4 environments for maize traits.

We then examined consistency between blocks and the contribution of block-related effects. For bean traits, correlations between blocks were mostly positive (144/153 positive; Table S5; Fig. S1–S4) with values ranging from 0.15 to 0.98 across environments. The block effect was significant for 47 of 56 bean traits and 21 of 30 maize traits, while the maize × block interaction was significant for 17 of 56 bean traits and 10 of 30 maize traits (Tables S6–S7). Bean and maize data were therefore adjusted using model M2 to account for block and maize × block interaction effects before subsequent analyses. Country and year effects were then assessed using two global mixed models (M3a and M3b) for maize and bean traits, respectively.

All maize traits showed strong G × E interactions (Table 2). Maize grown in Romania flowered later than in France (1034.19 °Cd vs. 881.89 °Cd in France), and produced markedly higher yields, with 47% higher thousand-kernel weight (TKW), 12% more ears per plant, and 33% greater kernel mass per plant. Most maize traits also varied among years. In 2022, maize flowered later (+6%) and was taller (+3%), whereas in 2023, plants produced 19% more ears, 20% more kernels, and 31% higher kernel mass per plant. Overall, maize growth was more vigorous in 2022, while yield was higher in 2023, particularly in Romania (Fig. S5).

**Table 2.**
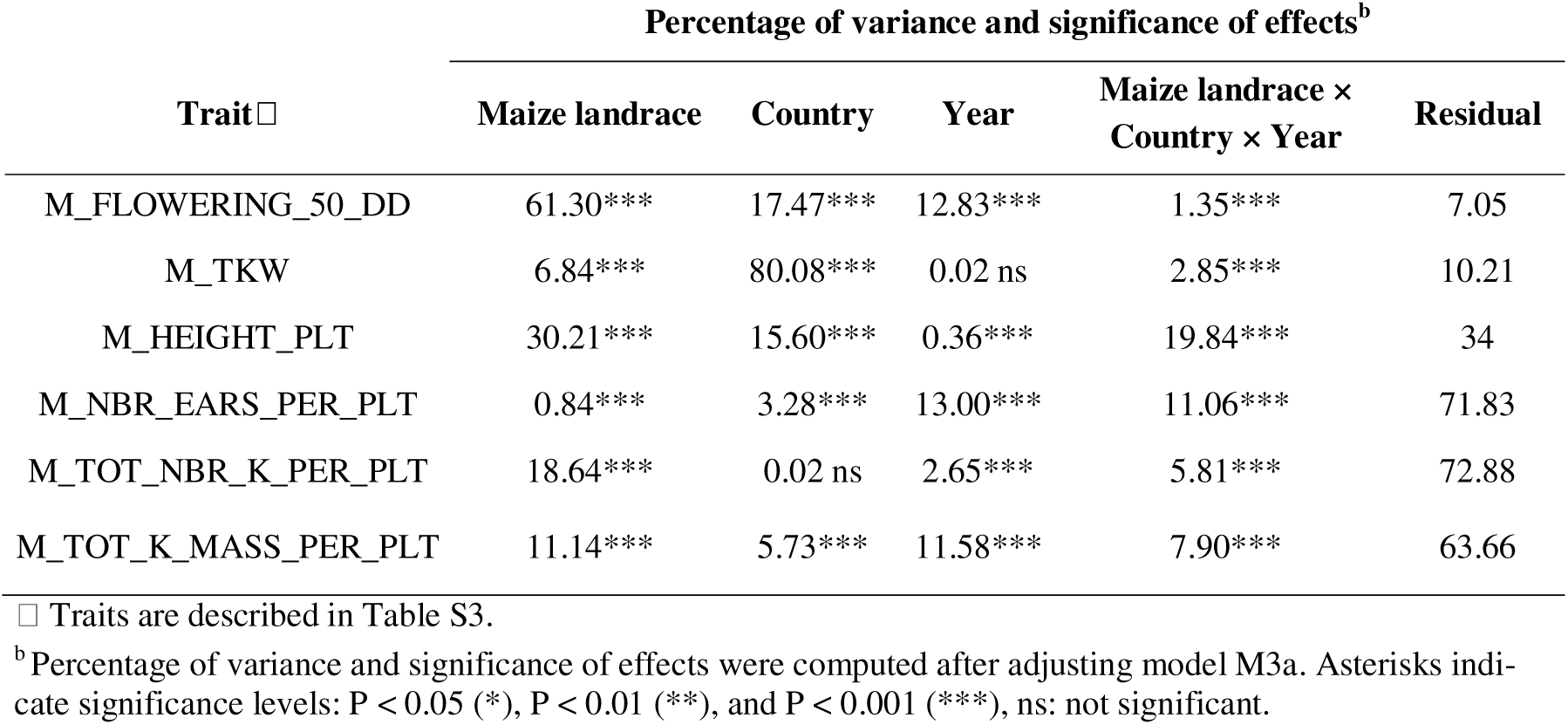
Fixed effects and percentage of variance for six maize traits.

| Trait□ | Percentage of variance and significance of effects<sup>b</sup> |  |  |  |  |
|---|---|---|---|---|---|
|  | Maize landrace | Country | Year | Maize landrace × Country × Year | Residual |
| M_FLOWERING_50_DD | 61.30*** | 17.47*** | 12.83*** | 1.35*** | 7.05 |
| M_TKW | 6.84*** | 80.08*** | 0.02 ns | 2.85*** | 10.21 |
| M_HEIGHT_PLT | 30.21*** | 15.60*** | 0.36*** | 19.84*** | 34 |
| M_NBR_EARS_PER_PLT | 0.84*** | 3.28*** | 13.00*** | 11.06*** | 71.83 |
| M_TOT_NBR_K_PER_PLT | 18.64*** | 0.02 ns | 2.65*** | 5.81*** | 72.88 |
| M_TOT_K_MASS_PER_PLT | 11.14*** | 5.73*** | 11.58*** | 7.90*** | 63.66 |

□ Traits are described in Table S3.
<sup>b</sup>Percentage of variance and significance of effects were computed after adjusting model M3a. Asterisks indi-cate significance levels: P < 0.05 (\*), P < 0.01 (\*\*), and P < 0.001 (\*\*\*), ns: not significant.

In bean, the country effect was predominant, being significant for 10 of the 12 measured traits, while the bean genotype × country interactions were significant for all traits (Table 3; Fig. S6). Phenology was markedly delayed in Romania, with flowering and pod formation occurring at 634 and 723 °Cd, respectively, compared to 440 and 462 °Cd in France. Yield and yield-related traits were generally higher in France, with 70% more pods per plant, 60% more seeds per plant, and 26% greater total seed mass per plant. Conversely, beans grown in Romania produced larger seeds, with 24% higher 100-seed mass and larger seed size (5% in length, 3% in width, 9% in surface area). Vegetative traits such as plant height and foliage density did not differ between countries. All bean traits, except thermal time to flowering, foliage density, and total seed mass per plant, varied between years (Table 3). Overall, beans performed better in 2023, with yield-related traits increasing by about 15% compared to 2022. The number of seeds per plant and per pod rose from 63 to 72 and from 3.7 to 4.0, respectively. In contrast, 100-seed mass was 11% lower in 2023 compared with 2022, consistent with slightly smaller seeds (-3% in surface area and -2% in width). Phenological timing remained stable across years, while plants were slightly shorter relative to maize in 2023 (2.05 vs. 2.16).

**Table 3.**
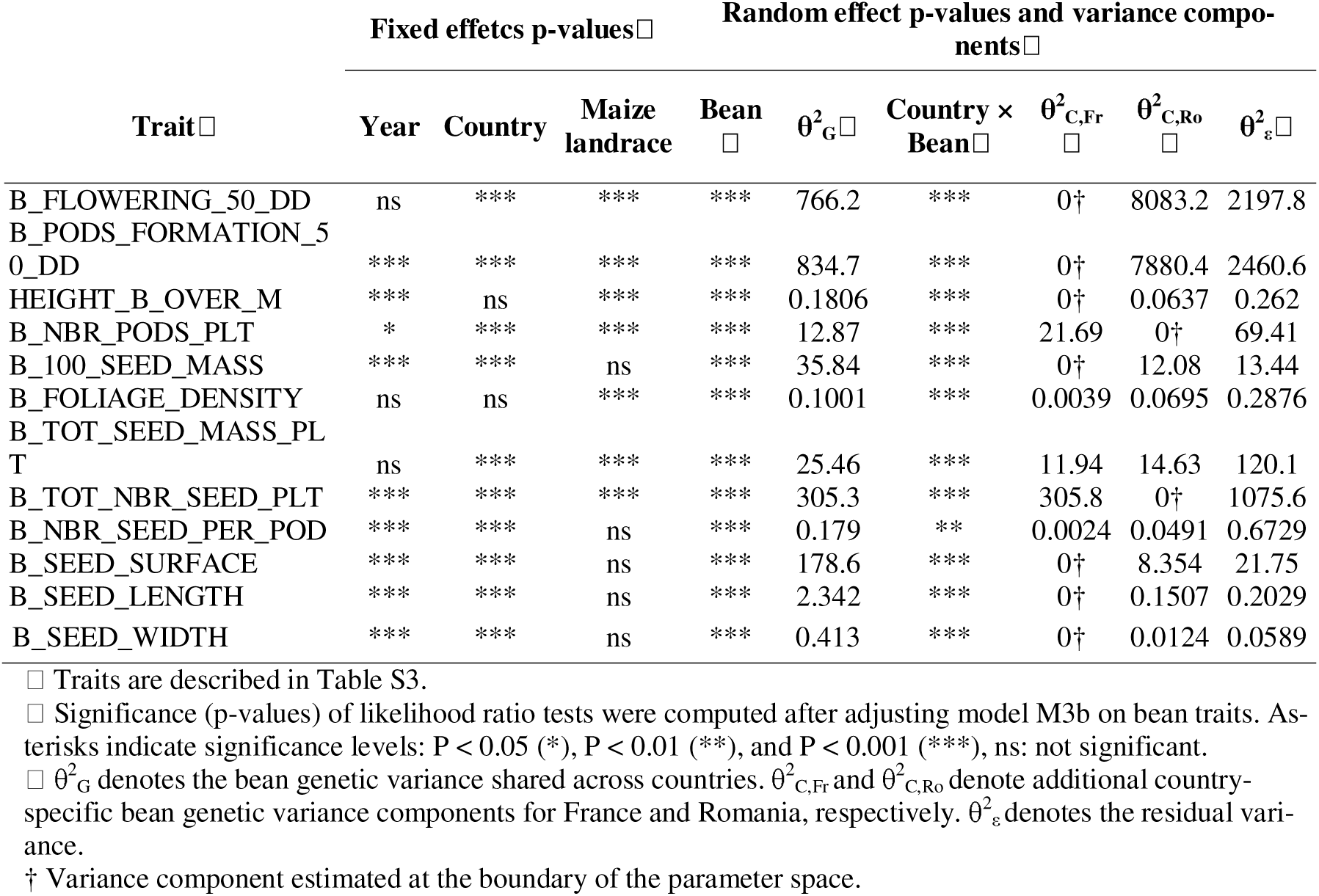
Statistical significance and variance components of the effects affecting bean traits.

| **Trait□** | **Fixed effetcs p-values□** |  |  | **Random effect p-values and variance compo-nents□** |  |  |  |  |  |
|---|---|---|---|---|---|---|---|---|---|
|  | **Year** | **Country** | **Maize landrace** | **Bean □** | **θ<sup>2</sup><sub>G</sub>□** | **Country × Bean□** | **θ<sup>2</sup><sub>C,Fr</sub> □** | **θ<sup>2</sup><sub>C,Ro</sub> □** | **θ<sup>2</sup><sub>ε</sub>□** |
| B_FLOWERING_50_DD | ns | *** | *** | *** | 766.2 | *** | 0† | 8083.2 | 2197.8 |
| B_PODS_FORMATION_50_DD | *** | *** | *** | *** | 834.7 | *** | 0† | 7880.4 | 2460.6 |
| HEIGHT_B_OVER_M | *** | ns | *** | *** | 0.1806 | *** | 0† | 0.0637 | 0.262 |
| B_NBR_PODS_PLT | * | *** | *** | *** | 12.87 | *** | 21.69 | 0† | 69.41 |
| B_100_SEED_MASS | *** | *** | ns | *** | 35.84 | *** | 0† | 12.08 | 13.44 |
| B_FOLIAGE_DENSITY | ns | ns | *** | *** | 0.1001 | *** | 0.0039 | 0.0695 | 0.2876 |
| B_TOT_SEED_MASS_PLT | ns | *** | *** | *** | 25.46 | *** | 11.94 | 14.63 | 120.1 |
| B_TOT_NBR_SEED_PLT | *** | *** | *** | *** | 305.3 | *** | 305.8 | 0† | 1075.6 |
| B_NBR_SEED_PER_POD | *** | *** | ns | *** | 0.179 | ** | 0.0024 | 0.0491 | 0.6729 |
| B_SEED_SURFACE | *** | *** | ns | *** | 178.6 | *** | 0† | 8.354 | 21.75 |
| B_SEED_LENGTH | *** | *** | ns | *** | 2.342 | *** | 0† | 0.1507 | 0.2029 |
| B_SEED_WIDTH | *** | *** | ns | *** | 0.413 | *** | 0† | 0.0124 | 0.0589 |

□ Traits are described in Table S3.
□ Significance (p-values) of likelihood ratio tests were computed after adjusting model M3b on bean traits. As-terisks indicate significance levels: P < 0.05 (\*), P < 0.01 (\*\*), and P < 0.001 (\*\*\*), ns: not significant.
□ θ<sup>2</sup><sub>G</sub> denotes the bean genetic variance shared across countries. θ<sup>2</sup><sub>C,Fr</sub> and θ<sup>2</sup><sub>C,Ro</sub> denote additional country-specific bean genetic variance components for France and Romania, respectively. θ<sup>2</sup><sub>ε</sub> denotes the residual vari-ance.
† Variance component estimated at the boundary of the parameter space.

Variance partitioning revealed a substantial genetic variance shared across countries for all traits, together with marked country-specific genetic variation. Additional genetic variance was predominantly observed in Romania, particularly for phenology and seed morphology, whereas variation in pod and seed number was mainly expressed in France; total seed mass showed additional genetic variance in both countries. Overall, all bean traits were influenced by bean genotype and genotype × country interactions, while maize landrace significantly affected 7 of the 12 traits, confirming strong environmental and G × E effects (Table 3). Since differences in mean values were greater between countries than between years (Fig. S6), maize × bean interactions were subsequently analyzed separately for each country. These analyses were performed after adjusting maize and bean data for year and maize × year effects using models M4a and M4b, respectively.

### 3.2. No evidence of local adaptation in beans

Most bean traits varied between countries and exhibited bean genotype × country interactions (Table 3). In addition, a latitudinal gradient was observed for traits measured in both France (SHM; Fig. S7a,b; Dim2 = 29.9% of total variance) and Romania (Bacău; Fig. S7c,d; Dim2 = 31.3%). Together, these patterns suggest that bean phenotypic variation may reflect local adaptation to the field sites. Using passport, genotypic, and climatic data for 183 bean lines, we tested whether performance in France (SMH) and Romania (Bacău) was related to the climatic distance between collection sites and experimental fields. Although PCA of 19 bioclimatic variables revealed strong climatic structuring among bean origins (Fig. S8), climatic distance was not associated with yield-related traits, providing no evidence for local adaptation. MRM further showed that genetic distance was the most frequent explanatory factor of phenotypic differentiation, followed by geographic distance, whereas bioclimatic distance had only marginal effects (Table S8).

### 3.3. The three maize landraces represent distinct biotic environments for beans

To assess the effects of each maize on bean development, phenology, and yield, we first characterized the three maize using data corrected for year effect and maize × year interaction (model M4a) in France and Romania, separately.

The three maize showed contrasting patterns of phenology, vegetative growth, and yield, with several significant country × maize interactions (Fig. 3; Fig. S9; Table 4). In both countries, most maize traits were positively correlated, except for thermal time to flowering, which was negatively correlated with most other traits (Fig. S10). Across both countries, Grand Roux Basque was the earliest maize, and produced shorter, narrower leaves, smaller ears (length and width), and lower kernel mass and number per plant, although it exhibited the highest thousand-kernel weight (TKW) and higher number of ears per plant. As a result, it was considered the least competitive maize. Gandino was consistently the latest-flowering maize, while Văleni showed intermediate phenology. In terms of vegetative growth, Gandino was the tallest in Romania, whereas Văleni was the tallest in France, both with similar leaf width. Regarding yield, Văleni achieved the highest TKW and number of ears per plant in France, and the highest total number of kernels per plant across both countries (Fig. 4; Fig. S9).

**Figure 3.**
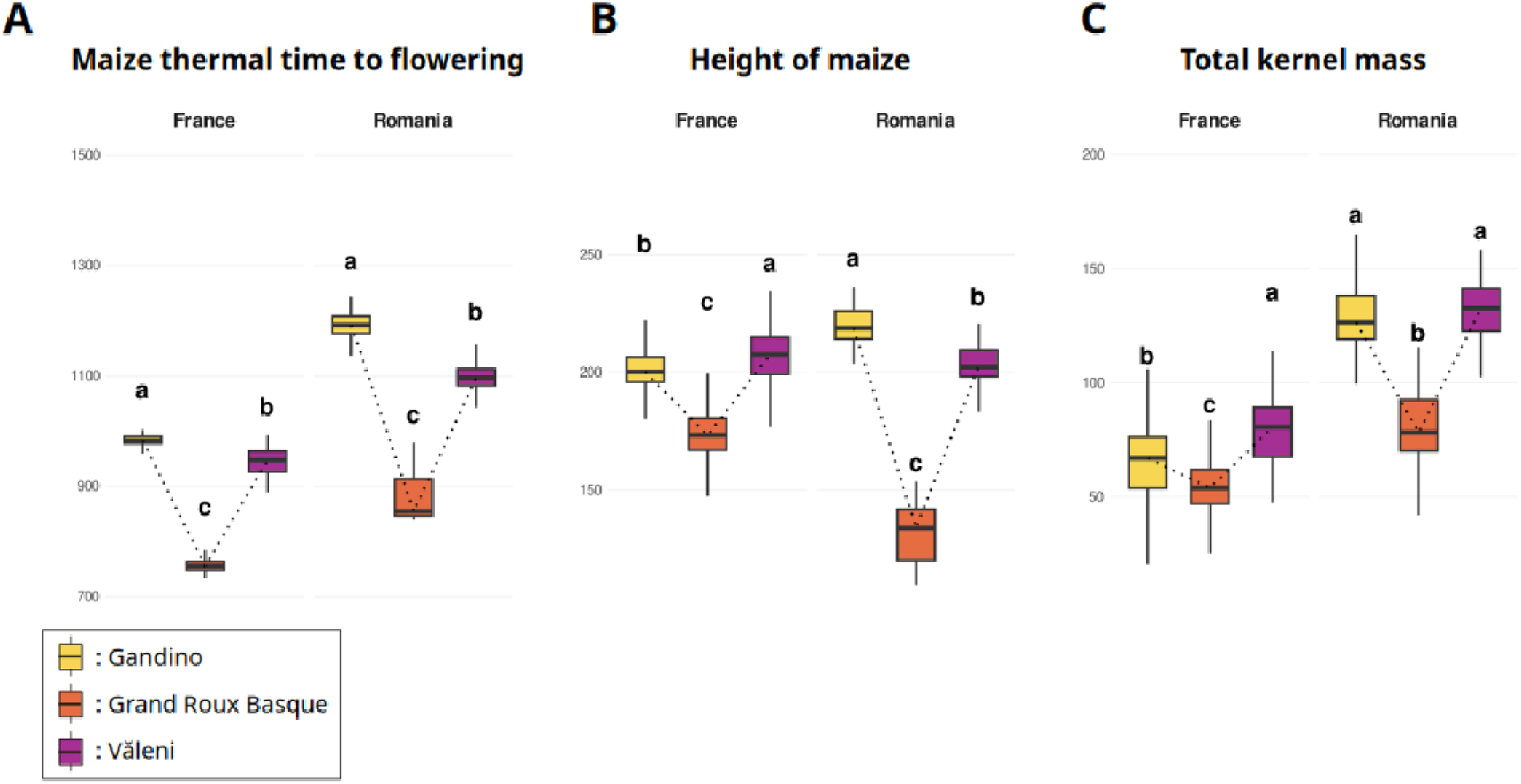
Boxplots of three maize traits measured in France and Romania for the three maize landraces. Traits represented include thermal time to flowering (M_FLOWERING_50_DD; A), plant height (M_HEIGHT_PLT; B), and total kernel mass per plant (M_TOT_K_MASS_PER_PLT; C). Values were corrected for block and year effects using model M4a. Different letters indicate statistically significant differences between maize landraces within each country based on Tukey-adjusted comparisons (P < 0.05). Sample sizes (n) correspond to the number of bean lines contributing to each boxplot. For thermal time to flowering (A), n = 201, 199, and 201 for Gandino, Grand Roux Basque, and Văleni, respectively, in France, and n = 43, 38, and 43, respectively, in Romania. For plant height (B), n = 201, 199, and 201 in France, and n = 43, 38, and 43 in Romania. For total kernel mass per plant (C), n = 201, 201, and 201 in France, and n = 43, 38, and 43 in Romania, for Gandino, Grand Roux Basque, and Văleni, respectively. ALT TEXT: Boxplots compare thermal time to flowering, height of maize, and kernel mass per plant among Gandino, Grand Roux Basque, and Văleni maize in France and Romania; letters indicate Tukey significance.

**Figure 4.**
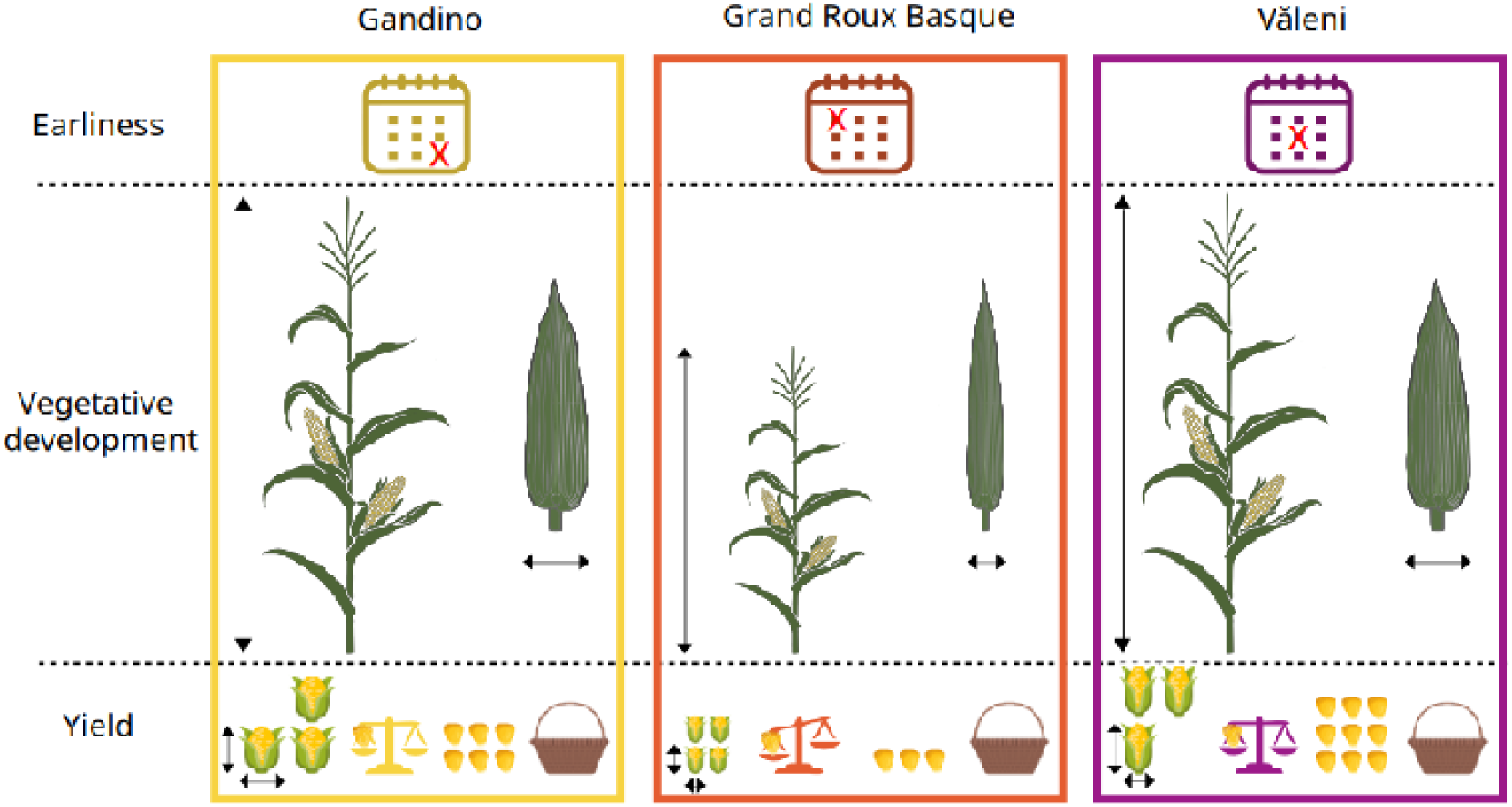
Summary of phenotypic differences among maize landraces. Traits are grouped into three categories: earliness, vegetative development, and yield. Each column represents a landrace, surrounded by a distinct border color. Thermal time to flowering is shown at the top (calendar icon), vegetative traits in the middle (plant height and leaf width), and yield traits at the bottom (from left to right: ear size and number per plant, thousand kernel weight, kernel number, and total kernel mass). Differences in size or number of icons/arrows reflects significant differences between landraces consistent across countries (France and Romania) using data corrected for block and year effects (using model M4a). ALT TEXT: Icon-based summary of maize landrace differences in flowering time, vegetative development, and yield traits that were consistent between France and Romania.

**Table 4.**
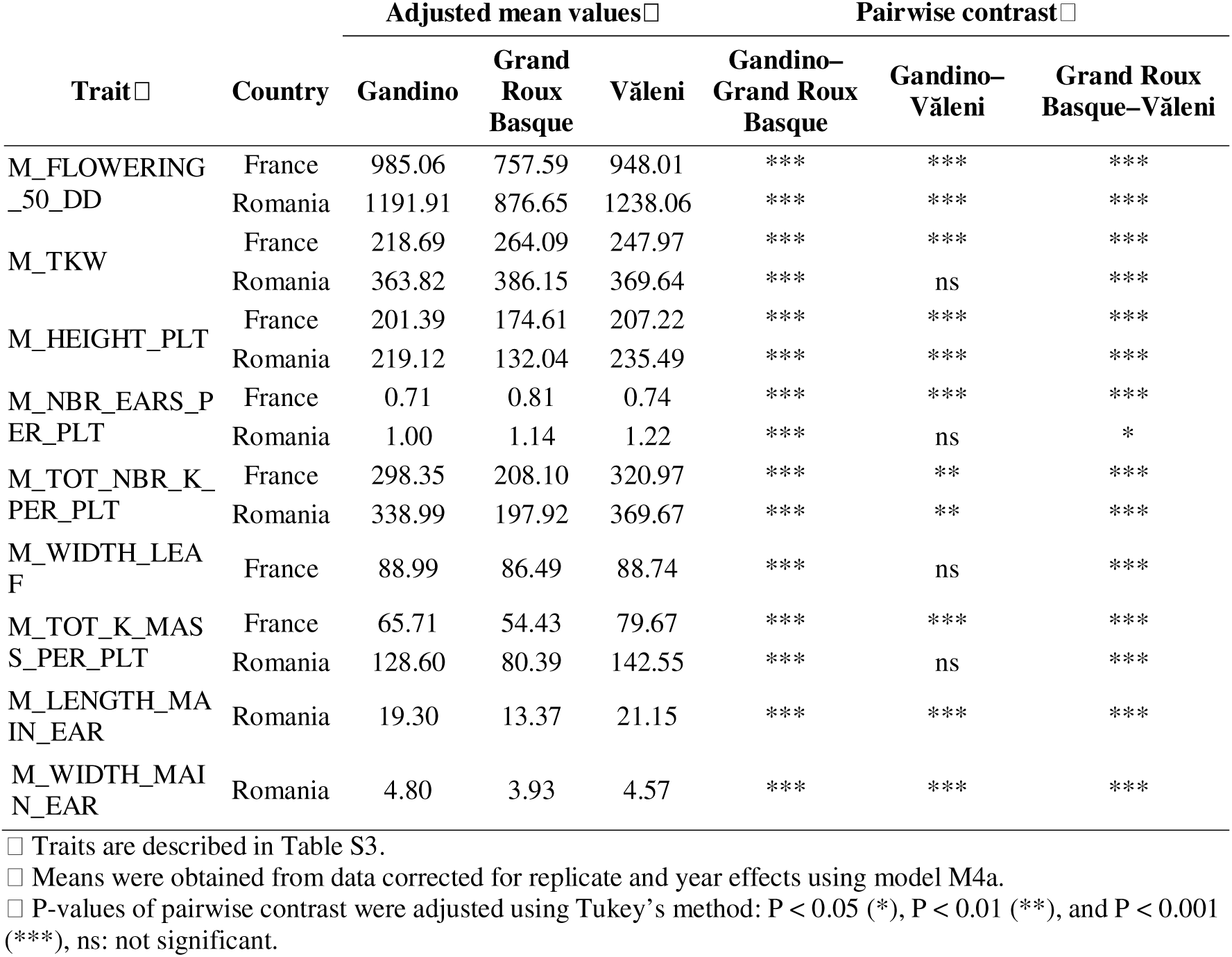
Mean trait values and differences among the three maize landraces in France and Romania.

|  |  | Adjusted mean values□ |  |  | Pairwise contrast□ |  |  |
|---|---|---|---|---|---|---|---|
| Trait□ | Country | Gandino | Grand Roux Basque | Văleni | Gandino–Grand Roux Basque | Gandino–Văleni | Grand Roux Basque–Văleni |
| M_FLOWERING_50_DD | France | 985.06 | 757.59 | 948.01 | *** | *** | *** |
|  | Romania | 1191.91 | 876.65 | 1238.06 | *** | *** | *** |
| M_TKW | France | 218.69 | 264.09 | 247.97 | *** | *** | *** |
|  | Romania | 363.82 | 386.15 | 369.64 | *** | ns | *** |
| M_HEIGHT_PLT | France | 201.39 | 174.61 | 207.22 | *** | *** | *** |
|  | Romania | 219.12 | 132.04 | 235.49 | *** | *** | *** |
| M_NBR_EARS_PER_PLT | France | 0.71 | 0.81 | 0.74 | *** | *** | *** |
|  | Romania | 1.00 | 1.14 | 1.22 | *** | ns | * |
| M_TOT_NBR_K_PER_PLT | France | 298.35 | 208.10 | 320.97 | *** | ** | *** |
|  | Romania | 338.99 | 197.92 | 369.67 | *** | ** | *** |
| M_WIDTH_LEAF | France | 88.99 | 86.49 | 88.74 | *** | ns | *** |
| M_TOT_K_MASS_PER_PLT | France | 65.71 | 54.43 | 79.67 | *** | *** | *** |
|  | Romania | 128.60 | 80.39 | 142.55 | *** | ns | *** |
| M_LENGTH_MAIN_EAR | Romania | 19.30 | 13.37 | 21.15 | *** | *** | *** |
| M_WIDTH_MAIN_EAR | Romania | 4.80 | 3.93 | 4.57 | *** | *** | *** |

□ Traits are described in Table S3.
□ Means were obtained from data corrected for replicate and year effects using model M4a.
□ P-values of pairwise contrast were adjusted using Tukey’s method: P < 0.05 (\*), P < 0.01 (\*\*), and P < 0.001 (\*\*\*), ns: not significant.

Multivariate analysis on phenotypic traits revealed distinct clustering of the three maize landraces, as illustrated in the French field (Fig. S11). They were clearly separated along the first PCA axis, which explained 44.4% of the total variance, with Grand Roux Basque being the most divergent, while Gandino and Văleni partially overlapped.

Maize also showed significant differences in variance for eight of the nine traits (Brown– Forsythe tests). Grand Roux Basque and Văleni showed the highest variance for most traits, whereas Gandino displayed the highest variance only for total kernel mass per plant (Table S9).

### 3.4. Above-ground phenotypic responses of bean to the three maize landraces

We compared bean phenotypes among maize for each country separately, using data corrected for year and year × maize effects (model M4b; Fig. 5; Fig. S12; Table S10). In both countries and across the three maize, most bean traits were positively correlated, except for seed size traits (surface, length, width, and 100-seed mass), which tended to show negative correlations with the number of pods per plant and the number of seeds per pod and per plant (Fig. S13). Overall, bean performance differed among maize, with some variation between countries (Fig. 6).

**Figure 5.**
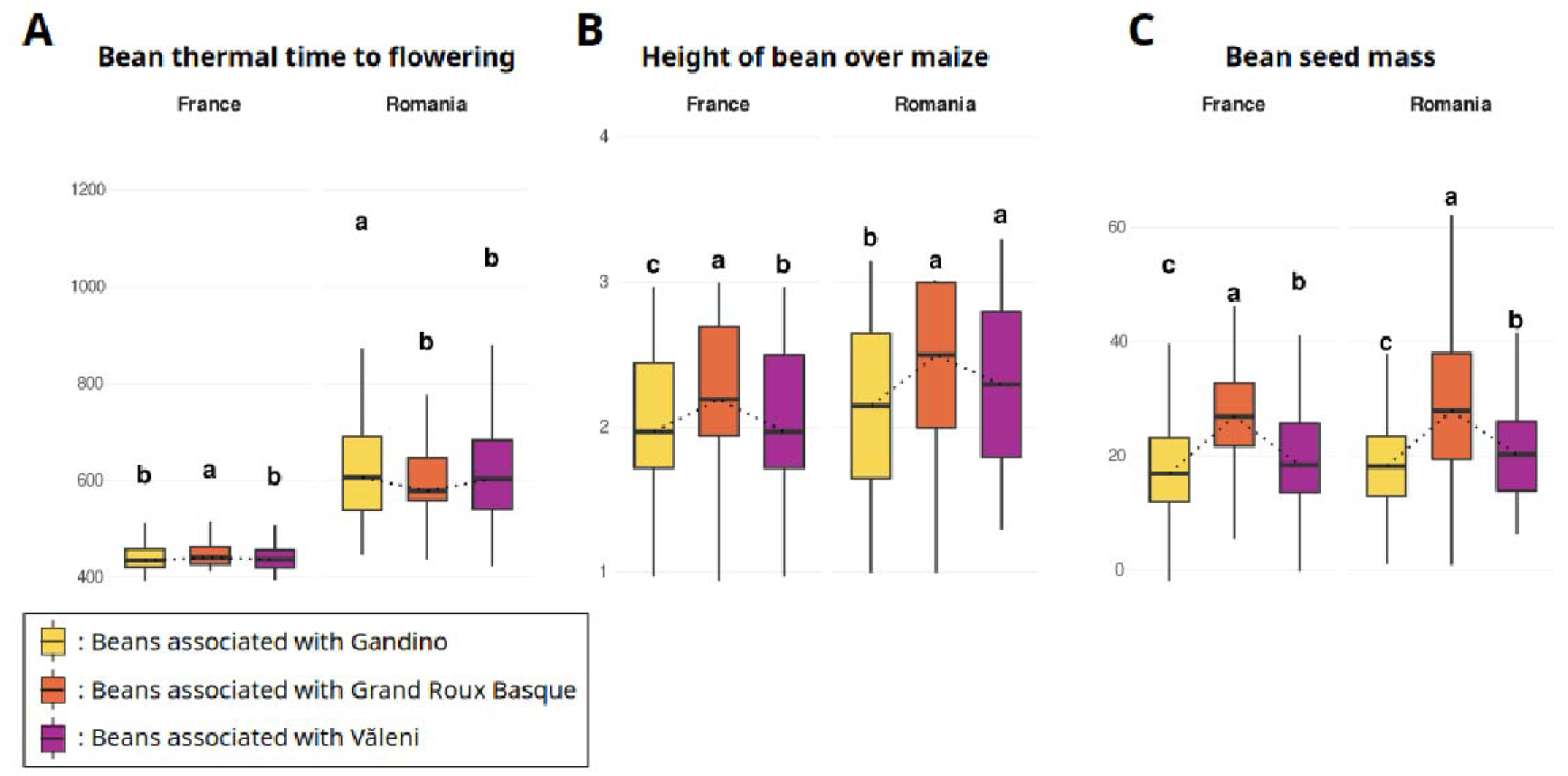
Boxplots of three bean traits measured in France and Romania. Traits represented include thermal time to flowering (B_FLOWERING_50_DD; A), relative bean plant height compared with associated maize (HEIGHT_B_OVER_M; B), and total seed mass per plant (B_TOT_SEED_MASS_PLT; C). Maize landrace environments are displayed in different colors. Values were corrected for block and year effects using model M4b. Different letters indicate statistically significant differences between maize landraces within each country. Sample sizes (n) correspond to the number of bean lines contributing to each boxplot. For thermal time to flowering (A), n = 199, 196, and 196 for Gandino, Grand Roux Basque, and Văleni, respectively, in France, and n = 199, 192, and 199, respectively, in Romania. For relative bean height (B), n = 201, 201, and 201 in France, and n = 200, 198, and 199 in Romania. For total seed mass per plant (C), n = 201, 201, and 201 in France, and n = 198, 193, and 198 in Romania, for Gandino, Grand Roux Basque, and Văleni, respectively. ALT TEXT: Boxplots compare bean flowering thermal time, bean height relative to maize, and seed mass per plant when bean lines were grown with each of the three maize landraces in France and Romania.

**Figure 6.**
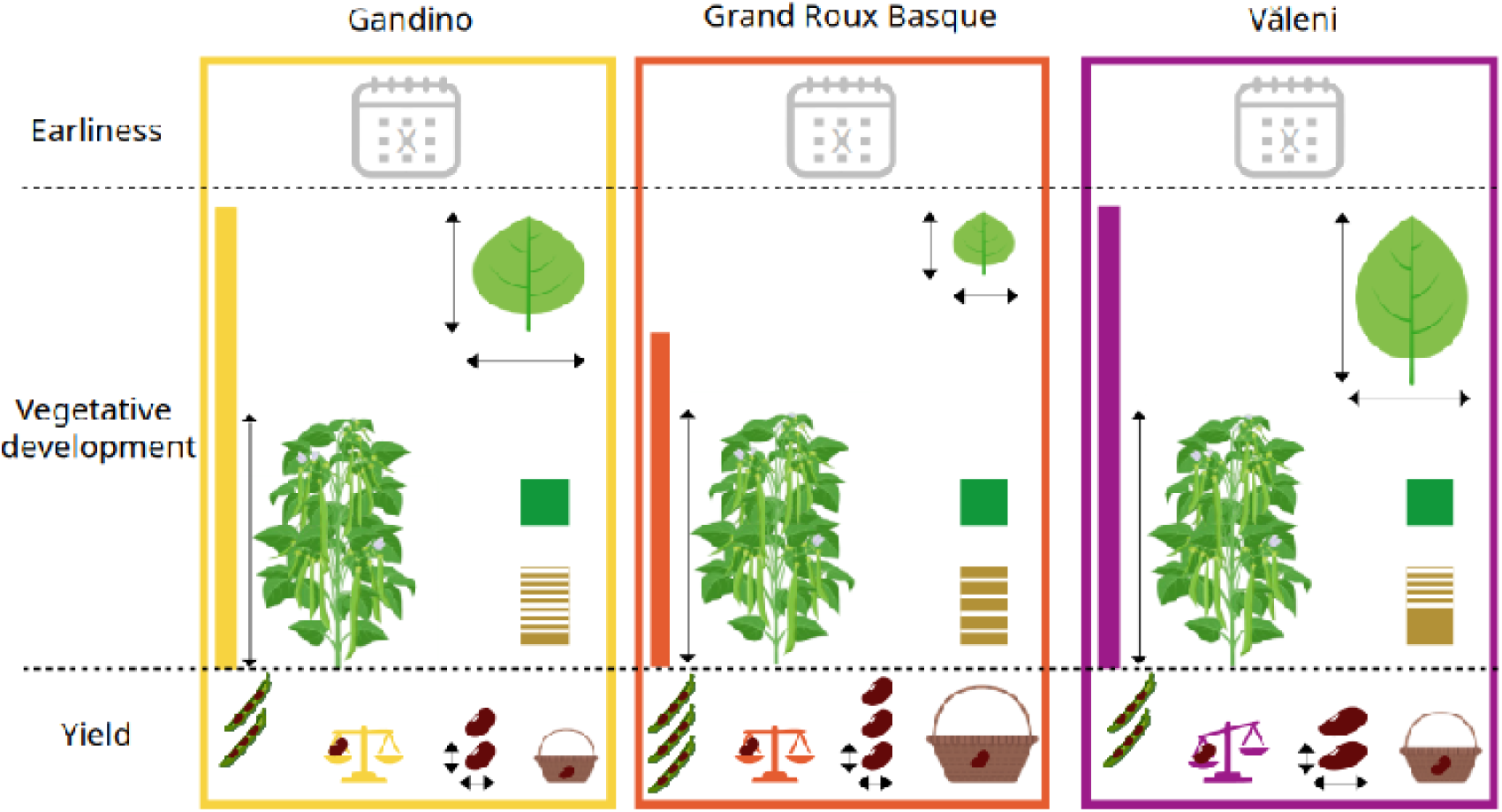
Summary of bean phenotypic differences among the three maize landraces. Traits are grouped in three categories: earliness, vegetative development, and yield. Each column corresponds to one maize landrace to which beans were associated, indicated by a distinct border color. Vegetative development is depicted with bean plant height relative to that of maize (maize being represented as a color bar and bean size as an arrow), size of the leaflets (length and width), foliage density (green squares), and pod placement (golden rectangles indicating pod placement along the bean plant). Yield, shown at the bottom, includes, from left to right, the numbers of pods per plant and seed per pod, the weight of 100 seeds, the number of seeds per plant, and the total seed mass per plant. Differences in size of arrows or icons, color or number of icons reflect the differences among maize landraces and their trend (larger/smaller). They were obtained from data corrected for block and year effects (using model M4b), and displayed only when consistent across countries (France and Romania). A description of traits is available in Table S3. ALT TEXT: Icon-based summary of bean differences among the three maize landrace environments for phenology, vegetative development, and yield traits that were consistent between countries.

In terms of phenology, beans intercropped with Gandino showed the latest flowering and pod formation in Romania, whereas ranks differed in France. For vegetative growth, beans associated with Grand Roux Basque consistently ranked highest for foliage density, relative height over maize, and homogeneous pod placement. In contrast, leaf surface was largest in beans intercropped with Văleni. Regarding yield, beans grown with Grand Roux Basque exhibited the highest total seed mass, number of pods, and number of seeds per plant, while beans associated with Văleni led to the highest 100-seed mass. Finally, beans consistently produced larger seeds (surface, length, and width) when intercropped with Văleni.

Overall, these patterns reflect the competitive hierarchy among maize: the least competitive, Grand Roux Basque, favored greater bean vegetative and reproductive performance. Its weak competitiveness, however, resulted in frequent lodging in Romania in 2022, which is why this maize landrace was not cultivated there in 2023.

### 3.5. Asymmetric competition structures maize–bean interactions

To determine whether the interaction between species was reciprocal or asymmetric, we examined the effects of each partner on the other, maize × bean interactions, cross-species trait correlations, and the consistency of high-performing combinations across environments.

Maize traits were largely unaffected by bean genotype (model M2; Table S7). In France, only maize thermal time to flowering and number of ears per plant (2022) and kernel mass per plant (2023) varied with bean genotype. Maize × bean interactions were rare and inconsistent across countries, being detected only for those same traits in 2022 in France, for kernel mass in 2023, and for ear number and ear width in Romania in 2022. In contrast, bean traits were strongly affected by the associated maize landrace, with this effect being significant for nearly all traits across both countries and years (Table S6). Maize × bean interaction occurred only in Romania in 2022 for traits related to bean earliness, vegetative growth, seed morphometry, and yield (Table S6).

We further explored maize × bean interactions by estimating Pearson’s correlation coefficients between maize and bean traits. Most significant correlations were negative, supporting competition between the two species. In France, 20 out of 25 maize-bean correlations were negative for Gandino, 15 out of 21 for Grand Roux Basque, and 21 out of 24 for Văleni. They were mostly observed between yield-related traits, with one exception: a positive correlation between kernel mass per plant of Grand Roux Basque and the 100-seed mass of its associated beans (Fig. S14). Positive correlations were generally specific to each maize, except for the correlation of maize flowering time and bean traits. Finally, bean phenology correlated positively with Gandino height but negatively with kernel weight and TKW of Văleni.

### 3.6. Interaction outcomes vary across maize partners and field experiments

To test whether interaction outcomes were consistent across maize partners and field experiments, we identified bean lines for which both bean and associated maize yields exceeded their respective medians within each field and maize landrace. These high-performing combinations were strongly maize-, year-, and country-specific, with overlap among groups no greater than expected by chance (Fig. 7). Historically co-cultivated bean lines were not enriched among them. In France, the balance-penalized performance index (MB_PERF) also differed among maize landraces, with Văleni highest overall, followed by Grand Roux Basque and Gandino (Fig. S15). Thus, interaction outcomes were highly context dependent: no single set of bean genotypes performed best across maize landrace and field experiments.

**Figure 7.**
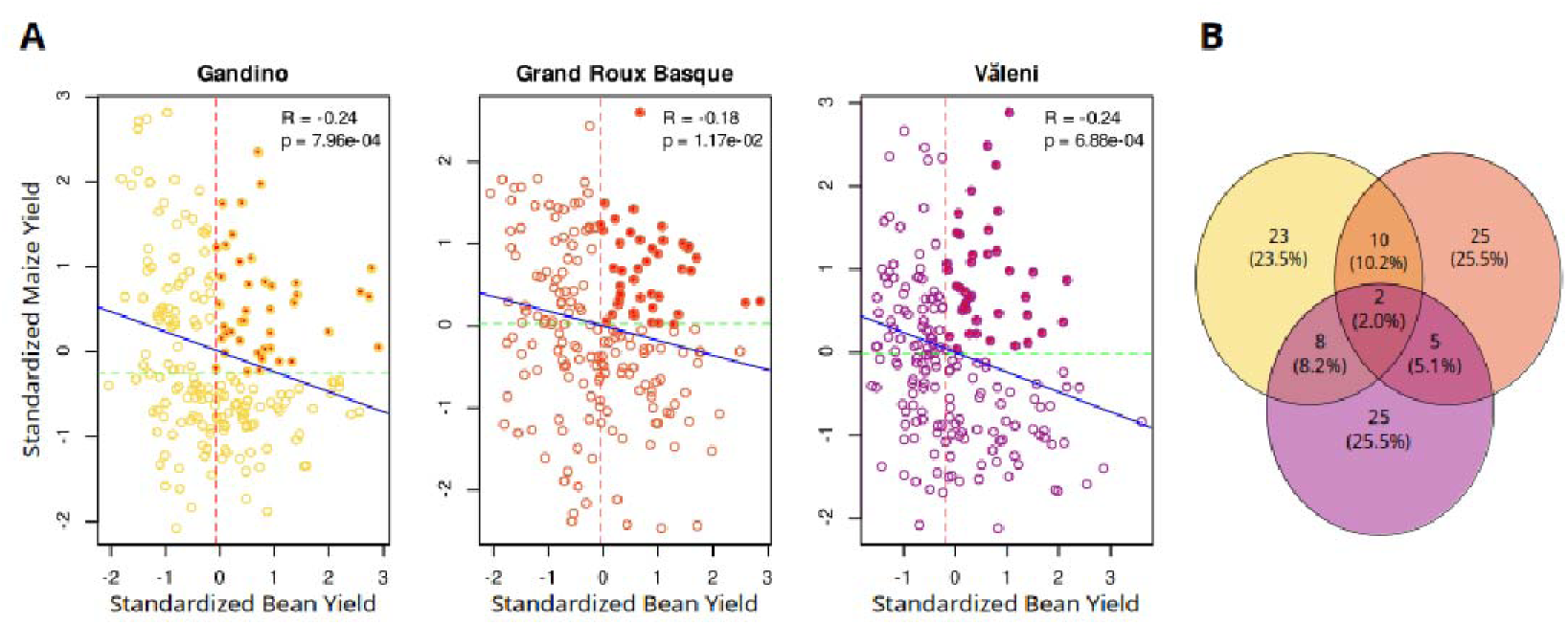
Identification of the best-performing maize-bean partners and overlap among maize landrace environments. Scatter plots show the relationship between standardized bean and maize yield (B_TOT_SEED_MASS_PLT and M_TOT_K_MASS_PLT for bean and maize, respectively) for each maize landrace (A). In the scatter plots, each dot represents a bean line. The Venn diagram illustrates the overlap among the best-performing bean lines (red dots in the scatterplots) in the three maize landrace contexts (B). Numbers (percentages) of bean lines are indicated. The figure is based on data from France corrected for block and year effects (using models M4a and M4b) and is standardized within each species and maize landrace. The blue lines represent the linear regression fits. The correlation coefficients (R) and the P-values (p) of the relationship are displayed in the top right of each plot. The dashed green and red lines indicate the bean and maize medians, respectively. Red dots highlight associations where the bean line has a standardized yield greater than the bean median yield, and the maize landrace has a standardized yield greater than the maize median yield. ALT TEXT: Three scatterplots relate standardized bean and maize yield for each maize landrace, with high-performing bean-maize combinations highlighted; a Venn diagram shows overlap of high-performing bean lines across maize backgrounds.

### 3.7. Below-ground maize × bean interactions

To assess below-ground interactions between maize and bean, we first characterized root phenotypes of the three maize landraces. Most of the variance in maize root traits measured manually did not depend on maize and/or year. Except for root angle (ANGLE), our model (M4a) explained less than 50% of the total variance in maize below-ground traits (Fig. S16). Root angle was the only trait showing a maize effect (Table S11, Fig. S16), but this effect varied among years (Fig. S16). In 2022, Văleni exhibited the smallest root angle, followed by Grand Roux Basque and Gandino, whereas in 2023 Gandino showed the smallest angle, followed by Văleni and Grand Roux Basque (Fig. S17).

In beans, root traits showed generally low mean heritability (Table S12), ranging from 0.075 to 0.41 for manually measured traits and from 0.033 to 0.457 for automatically measured traits. We discarded traits with heritability below 0.1 from further analyses (removing one manual and 20 automatic root traits; Table S12). The year effect had a major influence on all traits except nodulation score and basal angle (Table S13, Fig. S18, Fig. S19). The maize landrace affected five out of the nine manual traits and eight out of the 19 automatic traits, suggesting that the three associated maize represent distinct environments shaping bean below-ground architecture.

### 3.8. Stable and maize-dependent genetic effects underlie plant–plant interactions

Genotyping-by-sequencing and SNP imputation yielded 11,142 (MAF < 5%) and 3,004 (MAF < 25%) filtered SNPs for direct and indirect genetic-effect analyses, respectively. GWA conducted separately for each maize, country, and year were used to map bean traits (direct genetic effects) and maize traits (indirect genetic effects) onto the bean genome, followed by meta-GWA to distinguish effects that were stable across maize landraces and country–year combinations from effects that contrasted among maize landraces (Table S14, Fig. 8).

**Figure 8.**
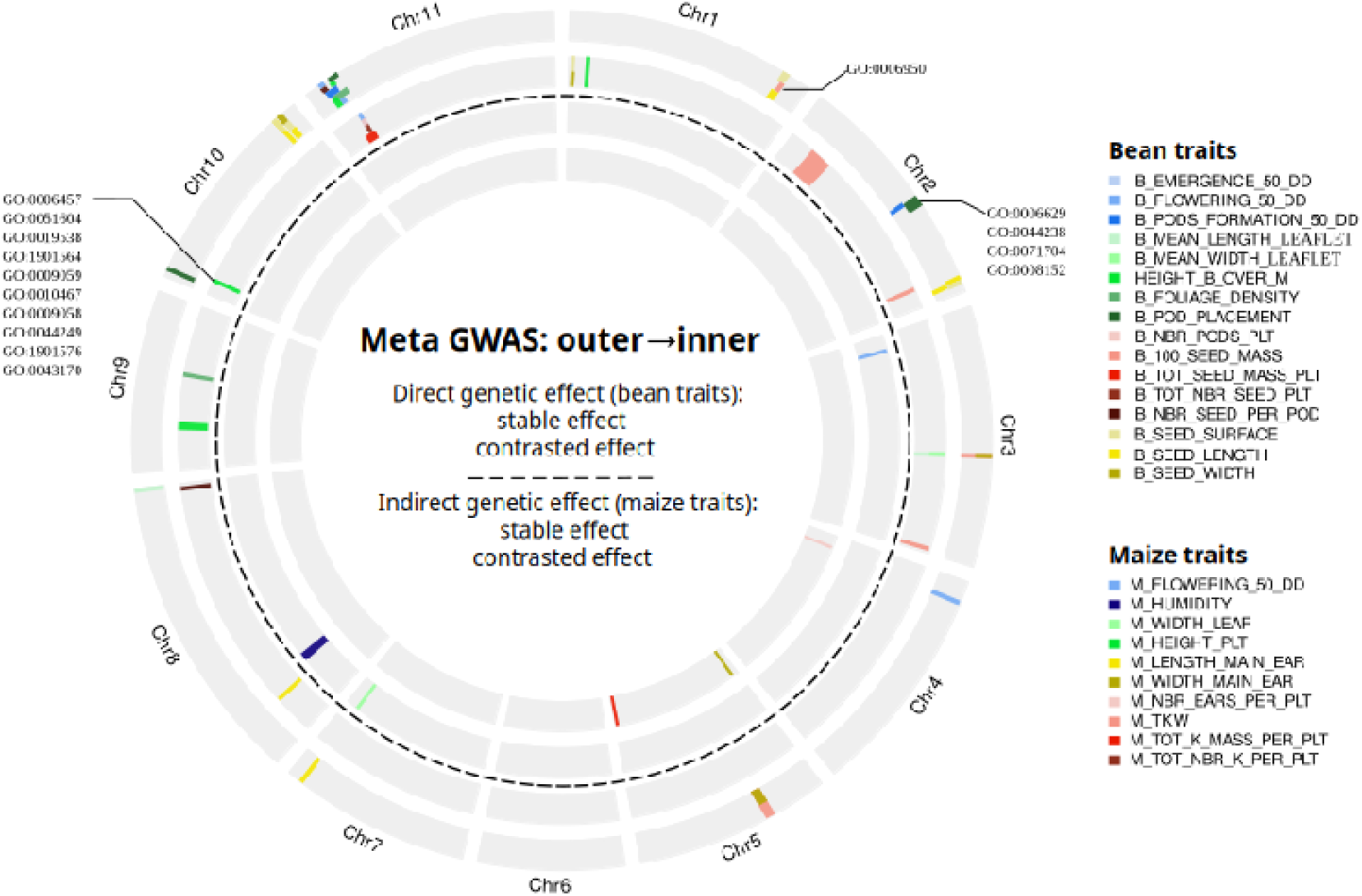
Genetic determinants of direct and indirect effects in maize–bean interactions as revealed by meta-GWA. Each of the four concentric circles represents the bean genome (11 chromosomes). Colored boxes indicate the physical positions of QTLs identified in each analysis. The two outer circles represent analyses for genetic determinants of direct genetic effects (involving bean traits) with stable and contrasting effects among maize landraces, and the two inner circles represent analyses for genetic determinants of indirect genetic effects (involving maize traits) with stable and contrasting effects among maize landraces. QTL colors correspond to traits indicated in the legend and described in Table S3. Enriched GO terms were associated with QTLs located on chromosome 1 (GO:0006950, response to stress), chromosome 2 (GO:0006629, lipid metabolic process; GO:0044238, primary metabolic process; GO:0071704, organic substance metabolic process; GO:0008152, metabolic process). QTL on chromosome 10 showed enrichment for multiple metabolic and biosynthetic processes, including protein folding (GO:0006457), protein maturation (GO:0051604), protein metabolic process (GO:0019538), organonitrogen compound metabolic process (GO:1901564), macromolecule biosynthetic process (GO:0009059), gene expression (GO:0010467), biosynthetic process (GO:0009058), cellular biosynthetic process (GO:0044249), organic substance biosynthetic process (GO:1901576), and macromolecule metabolic process (GO:0043170). ALT TEXT: Four concentric bean-genome rings summarize QTLs from stable- and contrasting-effect meta-GWA for direct bean effects and indirect effects on maize, with colors denoting traits and annotations marking Gene Ontology terms enrichment.

We identified 46 QTLs of direct genetic effects, among which 29 were stable, and 17 showed contrasting effects across maize. Eight stable direct effect QTLs, related to vegetative development and precocity, co-located on chromosome 11 with four QTLs with direct effects contrasted among maize, related to three yield traits and one phenological trait. Additionally, five direct-effect QTLs for seed morphometry formed two nearby clusters on chromosome 10, comprising two and three overlapping QTLs, respectively (Table S14, Fig. 8).

We identified only six bean QTLs with indirect-effect, among which three were stable across maize and three showed contrasting effects, with no overlap within or across analyses. Stable-effect QTLs were detected on chromosomes 3, 7, and 8, associated with maize flowering time, leaf width and kernel humidity, respectively. Contrasting-effect QTLs were located on chromosomes 4 (for ear number) and 5 (two independent regions, one associated with ear width and the other with kernel mass per plant).

GO analysis was performed on all candidate genes (Table S15) located within the identified QTL regions and revealed significant functional enrichment for the direct genetic effect analyses only. In the stable-effect meta-GWA, the QTL for 100-seed mass on chromosome 1 was enriched for response to stress, whereas the QTL for pod placement on chromosome 2 was enriched for terms related to lipid metabolic process, primary metabolic process, organic substance metabolic process, and metabolic process (Table S16). In the contrasted meta-GWA, the QTL for height of bean over the maize on chromosome 10 was enriched for ten GO terms such as protein folding and organonitrogen compound metabolic process (Table S16).

Combining GWA results from maize landrace-country-year combinations through meta-analysis increased statistical power and revealed QTLs that were not detected in individual analyses. More importantly, the contrast framework distinguished QTLs whose effects were robust to partner identity from those whose effects depended on the maize landrace.

In order to provide a more mechanistic interpretation of the results, we identified the orthologous genes in eight legume species and *Arabidopsis thaliana* corresponding to 2,043 unique *P. vulgaris* genes detected as candidate genes by meta-GWA (genes located within QTL intervals). The strongest candidates for plant-plant interactions were *Phvul.002G070900* and *Phvul.001G038400*, together with the nearby *Phvul.001G038800* (*CKX7*). Eleven genes corresponded to loci associated with indirect genetic effects in *Arabidopsis thaliana*, including *Phvul.007G243200*, which overlapped the stable indirect-effect QTL for maize leaf width. Additional candidates were associated with jasmonate signaling, peptide transport, nitrogen acquisition, and plant architecture (Supplementary Tables S17 and S18).

## 4. Discussion

Although intercropping has long been promoted as an ecological strategy to enhance agricultural sustainability (Vandermeer, 1995; Malézieux *et al*., 2009; Bedoussac *et al*., 2015; Li *et al*., 2023), the genetic bases of plant–plant interactions remain poorly understood. Here, we show that interspecific interactions between common bean and maize are asymmetric and environmentally contingent. Bean responses are shaped by maize partner identity and indirect genetic effects are either stable or partner-dependent. These patterns emerged across a diverse set of 200 bean lines, with no evidence that bean responses were driven by climatic local adaptation. Phenotypic differentiation among bean lines was instead associated primarily with genetic and geographic distances, consistent with previous work highlighting the role of historical and cultural factors shaping European bean diversity (Bellucci *et al*., 2023).

### 4.1. Competitive asymmetry shapes plant responses to the biotic environment

Cereals are commonly the dominant competitors in cereal–legume mixtures (Ofori and Stern, 1987; Baudoin *et al*., 1997; Corre-Hellou *et al*., 2006; Lithourgidis *et al*., 2011; Annicchiarico *et al*., 2017). Our results are consistent with this well-established asymmetry: maize acted as the dominant species, with several maize traits negatively affecting bean yield-related traits, while bean exerted comparatively a negligible influence on maize performance (Table S9). This pattern was also observed in a previous study showing that maize–bean intercropping strongly altered the bean transcriptomic response, whereas the maize transcriptome remained largely unaffected (Vazeux-Blumental *et al*., 2025). Under such asymmetric interactions, variation among bean lines can be interpreted primarily as variation in competitive tolerance—the ability to maintain development and reproduction in the presence of a dominant neighbor, consistent with the findings of Davis and Garcia (1983).

Bean lines combining high bean and maize yields showed limited overlap across maize partners and field environments, and historically co-cultivated bean lines were not overrepresented among these high-performing combinations. In contrast, the highest mixture performances were observed with Văleni, the only landrace in a documented history of intercropping in Transylvania (>50 years). Although the specific traits underlying its superior performance remain unknown, its longstanding co-cultivation with bean likely contributed to its suitability for intercropping.

### 4.2. Above- and below-ground phenotypic responses to genetically distinct neighbors

The effects of neighbor identity extended both above and below-ground traits. Above ground, beans differed among maize partners in relative height, foliage development, phenology, reproductive allocation, and seed morphology, consistent with responses to the competitive environment created by neighboring plants. Below-ground responses were more difficult to characterize. Although maize and bean roots were consistently intertwined under field conditions, the effect of the maize landrace identity on the phenotypic variation of maize root traits was generally limited (Fig. S16). Bean root traits actually showed very low heritability (Table S12), which may partly reflect the limited sample size, given that Strock *et al*. (2019) reported heritabilities above 0.52 for similar manually measured root traits using a much larger dataset of 577 accessions evaluated across 17 environments and five years. Nevertheless, maize landrace identity affected several architectural traits of bean roots such as the number of basal root whorls and the basal root angle, although these effects varied substantially between years (Table S13). This indicates that the below-ground influence of maize on bean root development is shaped by environmental conditions. Previous controlled studies have shown that root plasticity in intercropping can improve access to limiting resources, including water, nitrogen, and phosphorus, through changes in root distribution and rhizosphere processes (Li et al., 2006; Zhang et al., 2022). Our results extend this perspective to field-grown maize–bean intercrops, highlighting both the potential for neighbor-dependent root responses and the challenges of detecting them under field conditions.

### 4.3. Meta-GWA reveals stable and maize-landrace-specific genetic effects

Meta-analysis of 11 GWAs increased statistical power and enabled detection of G × E interactions through contrasts among maize landraces, extending combining ability concepts from hybrid breeding (Sprague and Tatum, 1942). We identified QTLs with stable effects across landraces, analogous to General Mixing Ability (GMA), and QTLs with landrace-specific effects, analogous to Specific Mixing Ability (SMA), which captures pair-specific interactions (Jensen and Federer, 1965). We detected 46 QTLs for direct genetic effects on bean traits, whereas only six QTLs were associated with maize traits. This imbalance may largely result from lower statistical power in the indirect-effect analysis, due to the smaller number of phenotyped maize plants, fewer markers, and the inherently weaker genetic control of neighbor effects. Several QTLs were detected for bean phenology, vegetative development, yield components, and seed morphology. Few colocalized with major common bean genes such as *PvTFL1y/fin* (growth habit) or *PHYA3/Ppd* (photoperiod response and flowering time) (Repinski *et al*., 2012; Weller *et al*., 2019). One notable candidate, *Phvul.010G130600* within the stable seed-length QTL on chromosome 10, corresponds to the J locus associated with seed coat darkening (Erfatpour and Pauls, 2020). Although it is not known as a causal gene for seed length, its presence in a seed morphology QTL may indicate linkage with broader seed-related variation, or alternatively a pleiotropic effect on multiple seed traits. This is consistent with evidence in pea showing that domestication altered seed coat gene expression and secondary metabolite composition, suggesting that seed coat-related loci may affect seed characteristics (Klčová *et al*., 2024).

### 4.4. Candidate genes point towards neighbor perception and resource acquisition

Identifying genes underlying intercropping performance requires integrating above- and belowground biological processes associated with competition, neighbor responses, nutrient acquisition, hormonal regulation, nitrogen-fixing symbioses, and plant architecture. We conducted orthologous search (Table S17) and a literature survey (Table S18). The strongest candidates were *Phvul.002G070900* and *Phvul.001G038400*, both conserved across the legume species examined. *Phvul.002G070900* is orthologous to the *Arabidopsis thaliana* receptor kinase *ESCAPE1* (PERK13), recently identified as a regulator of neighbor-induced competitive responses and validated through mutant analysis (Libourel *et al*., 2026). *Phvul.001G038400* corresponds to the Arabidopsis *thaliana ERECTA* gene, a regulator of plant architecture whose natural allelic variation is associated with reduced individual competitiveness but increased productivity at the group level (Biernaskie *et al*., 2025). The adjacent gene *Phvul.001G038800*, encoding cytokinin oxidase/dehydrogenase 7 (*CKX7*), is an additional candidate because cytokinins influence root and shoot growth, architecture and, nutrient acquisition function (Köllmer *et al*., 2014), and the corresponding locus has been associated with competitive performance in chickpea (Lake *et al*., 2016).

A second group of 11 candidates was identified through orthology with *Arabidopsis thaliana* loci controlling indirect genetic effects (Montazeaud *et al*., 2023). Most occurred within QTL intervals controlling direct genetic effects on bean traits including seed size, foliage density and pod placement (Table S15). *Phvul.007G243200*, encoding an ABC transporter-related protein, was uniquely located within a stable indirect-effect QTL affecting maize leaf width in maize-bean mixtures (Table S15). This makes it one of the strongest candidates for genotype-dependent effects of common bean on its cereal neighbor. Several additional candidates were supported by transcriptomic studies of cereal–legume intercrops. These include *MYC2*-family transcription factors (*Phvul.003G285700* and *Phvul.002G141500*), which are central regulators of jasmonate-mediated signaling (Kazan and Manners, 2013) and a chloroplastic linoleate 13S-lipoxygenase 2-1 (*Phvul.011G056500*) which participates to jasmonate biosynthesis; a *PTR1*-like transporter (*Phvul.008G107200*), which is involved in peptide uptake and root responses (Komarova *et al*., 2008), and exhibits a higher expression in peanut roots under peanut-maize intercropping (Dai *et al*., 2018).

Several additional candidate genes may contribute to architectural and symbiotic responses under intercropping. *Phvul.003G291500*, an orthologue of the Arabidopsis GAI DELLA protein, regulates plant architecture and belowground responses through gibberellin signaling, consistent with evidence linking gibberellin-mediated pathways to nodulation and nitrogen fixation in maize–pea intercropping (Peng *et al*., 1997; Sun *et al*., 2026). *Phvul.010G129400*, encoding an NSP1-like GRAS transcription factor, is a strong candidate for genotype-dependent variation in nodulation and microbial interactions because of the established role of NSP1 proteins in symbiotic signaling (Smit *et al*., 2005; Yu *et al*., 2026). *Phvul.011G045900*, orthologous to Arabidopsis AGB1, influences carbon allocation and root system plasticity under plant competition through its role in sugar and auxin signalling (Mudgil *et al*., 2016).

These findings have implications beyond intercropping. They highlight that a plant’s phenotype is shaped not only by its own genotype and the abiotic environment, but also by the genetic identity of neighboring plants. By demonstrating that genetic variation can influence the phenotype of neighboring individuals through indirect genetic effects, and that these effects depend on environmental context, our study contributes to a broader understanding of plant-plant interactions. Considering the genotypes of neighboring plants as components of the biotic environment provides a framework for integrating quantitative genetics, phenotypic plasticity, and the molecular mechanisms underlying plant–plant interactions. This approach is relevant to both managed and natural plant communities.

## Supplementary data

Figure S1. Pearson correlations between blocks at the French field site in 2022.

Figure S2. Pearson correlations between blocks at the French field site in 2023.

Figure S3. Pearson correlations between blocks at the Romanian field site in 2022.

Figure S4. Pearson correlations between blocks of raw data at the Romanian field site in 2023.

Figure S5. Distributions of maize traits across all experimental fields.

Figure S6. Distributions of bean traits across all experimental fields.

Figure S7. Bean distribution shows a latitudinal gradient.

Figure S8. PCA of 183 European bean lines based on 19 bioclimatic variables.

Figure S9. Boxplots of nine maize traits measured in France and/or Romania.

Figure S10. Correlation matrices between maize traits.

Figure S11. PCA of the three maize landraces.

Figure S12. Boxplots of fifteen bean traits measured.

Figure S13. Pearson’s correlation matrices between bean traits.

Figure S14. Pearson correlation matrices between maize and bean traits.

Figure S15. Distribution of the cumulated maize and bean yield.

Figure S16. Variance decomposition for the seven maize below-ground traits.

Figure S17. Photos illustrating the three maize landrace root systems.

Figure S18. Boxplots of eight manually measured below-ground bean traits.

Figure S19. Boxplots of 19 automatically measured below-ground bean traits.

Table S1. Passport data of the 201 bean accessions and three maize varieties included in the study.

Table S2. Initial bean panel and admixture coefficients.

Table S3. Maize, bean, and common traits measured above- and below-ground.

Table S4. Description of 19 bioclimatic variables extracted from the CHELSA V2.1 database.

Table S5. Pearson correlations between blocks for bean traits.

Table S6. P-values of likelihood ratio tests after adjusting model M2b on bean traits.

Table S7. P-values of likelihood ratio tests after adjusting model M2a on maize traits.

Table S8. Bean local adaptation evaluated with multiple regression on distance matrices.

Table S9. Variance within the three maize landraces.

Table S10. Differences of bean response to maize.

Table S11. P-values from the ANOVA model M4a adjusted on maize root traits.

Table S12. Mean broad-sense heritability of bean root traits.

Table S13. P-values of likelihood ratio tests after adjusting model M4b on bean root traits.

Table S14. QTL obtained from four meta-GWA analyses.

Table S15. Candidate genes found within QTL intervals for the four meta-GWA analyses.

Table S16. GO terms of genes located within bean QTLs detected in the four meta-GWA analyses.

Table S17. Orthologue and OrthoFinder orthogroup assignments for QTL-associated *Phaseolus vulgaris* candidate genes across the analysed species.

Table S18. Comparative evidence supporting prioritization of QTL-associated *Phaseolus vulgaris* candidate genes implicated in plant-plant interactions and cereal-legume intercropping.

## Acknowledgements

We warmly thank Aline Supper, Jean-René Loustalot, Angéline Pourtau, Arnaud Brunet, Silvica Ambăru□, Andreea Antal Tremurici, Mariana Calara, Dan Ioan Avasiloaiei, Alin Gabiel Iosob, Claudia Bălăi□ă, Arthur Wojcik, Harry Belcram, Fabrice Dumas, Ester Murube, and Tania Gioia for their valuable contribution and support throughout this work.

ChatGPT (OpenAI) was used between 2 September 2026 at 16:26 CEST and 12 October 2026 at 14:30 CEST, with prompts requesting verification of the manuscript against the Journal of Experimental Botany author guidelines, assistance with language editing, and preparation of alternative text descriptions for figures. All suggested changes were reviewed and validated by the authors, who take full responsibility for the final content.

## Author contributions

Noa Vazeux-Blumental: Methodology, Software, Validation, Formal analysis, Investigation, Data curation, Writing — original draft, Visualization, Funding acquisition. Carine Palaffre: Investigation, Resources. Creola Brezeanu: Investigation, Resources. Petre M. Brezeanu: Investigation. Bernard Lagardère: Investigation. Cyril Bauland: Investigation, Resources, Writing — review & editing. James Burridge: Methodology, Investigation, Writing — review & editing. Pablo Affortit: Investigation. Alexandre Grondin: Methodology, Investigation, Writing — review & editing. Marzia Rossato: Methodology, Software, Resources, Writing — review & editing. Andrea Benazzo: Methodology, Software, Resources. Tristan Mary-Huard: Methodology, Writing — review & editing. Laurence Moreau: Methodology, Writing — review & editing. Roberto Papa: Writing — review & editing, Project administration, Funding acquisition. Elisa Bellucci: Methodology, Resources, Writing — review & editing. Elena Bitocchi: Methodology, Validation, Resources, Writing — review & editing. Filippo Servalli: Resources. Laurent Laplaze: Methodology, Investigation, Writing — review & editing. Domenica Manicacci: Conceptualization, Methodology, Investigation, Writing — review & editing, Supervision, Funding acquisition. Maud I. Tenaillon: Conceptualization, Methodology, Resources, Writing — review & editing, Supervision, Project administration, Funding acquisition.

## Data availability

Phenotypic, bioclimatic, and genomic data are available in Recherche Data Gouv (https://doi.org/10.57745/HACRRX).

## Abbreviations

FDR: false discovery rate
GBS: genotyping-by-sequencing
GO: Gene Ontology
GWA: genome-wide association
HSD: honestly significant difference
LD: linkage disequilibrium
LS: local score
MAF: minor allele frequency
MRM: multiple regression on distance matrices
PCA: principal component analysis
QTL: quantitative trait locus
SMH: Saint-Martin-de-Hinx
SNP: single nucleotide polymorphism
SSD: single seed descent
TKW: thousand-kernel weight.

