## Supplementary figures for "Genetic effects of plant–plant interactions depend on neighbor identity and environmental context"

**Journal of Experimental Botany**

**Noa Vazeux-Blumental et al.**

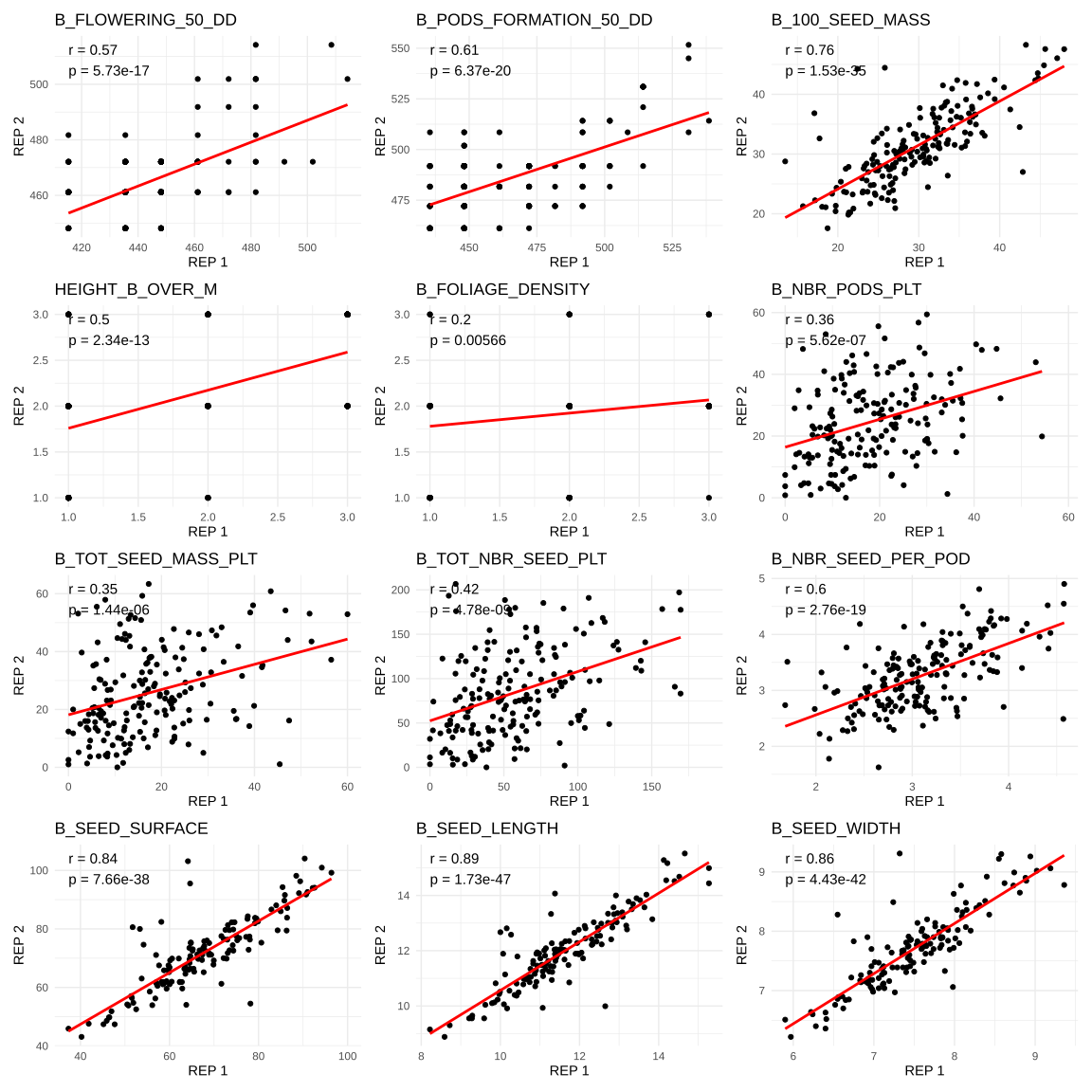

Figure S1. Pearson correlations between block 1 (REP 1) and block 2 (REP 2) of raw data for 12 bean traits (as defined in Table S3) grown in association with the Gandino maize at the French field site in 2022. Red lines represent the linear regression of block 2 on block 1. For each trait, the correlation coefficient (r) and *P*-value (p) are displayed at the top-left of the plot.

**ALT TEXT:** Twelve scatterplots compare block 1 and block 2 measurements for bean traits in France in 2022 with Gandino maize. Red regression lines show generally positive block-to-block relationships, strongest for seed size and morphology traits and weaker for several developmental traits; each panel reports Pearson r and P.

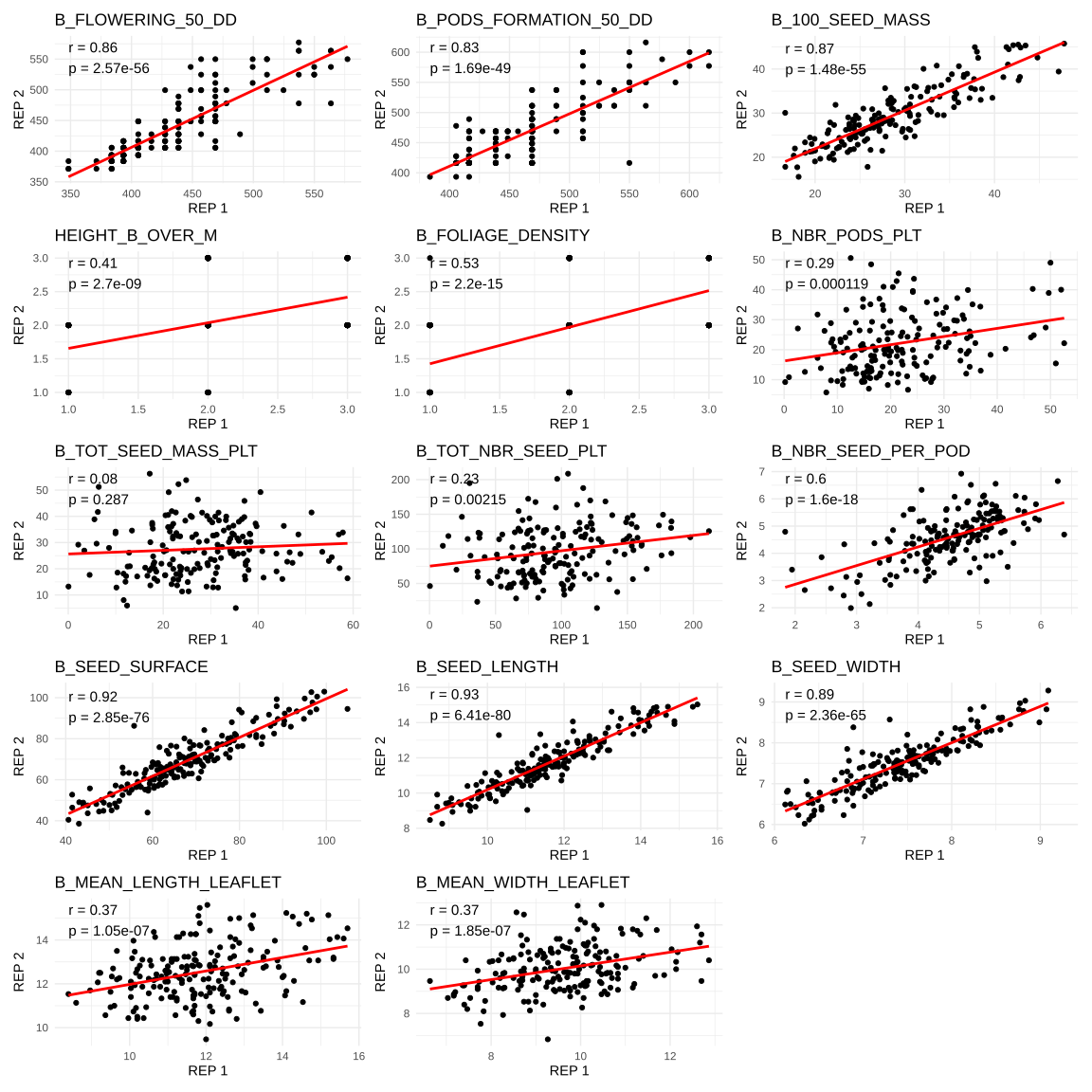

Figure S2. Pearson correlations between block 1 (REP 1) and block 2 (REP 2) of raw data for 14 bean traits (as defined in Table S3) for plants grown in association with the Gandino maize at the French field site in 2023. Red lines represent the linear regression of block 2 on block 1. For each trait, the correlation coefficient (r) and *P*-value (p) are displayed at the top-left of the plot.

**ALT TEXT:** Fourteen scatterplots compare block 1 and block 2 measurements for bean traits in France in 2023 with Gandino maize. Most seed size and morphology traits show tight positive relationships between blocks, whereas some phenology, yield, and foliage traits are more dispersed; each panel reports Pearson r and P.

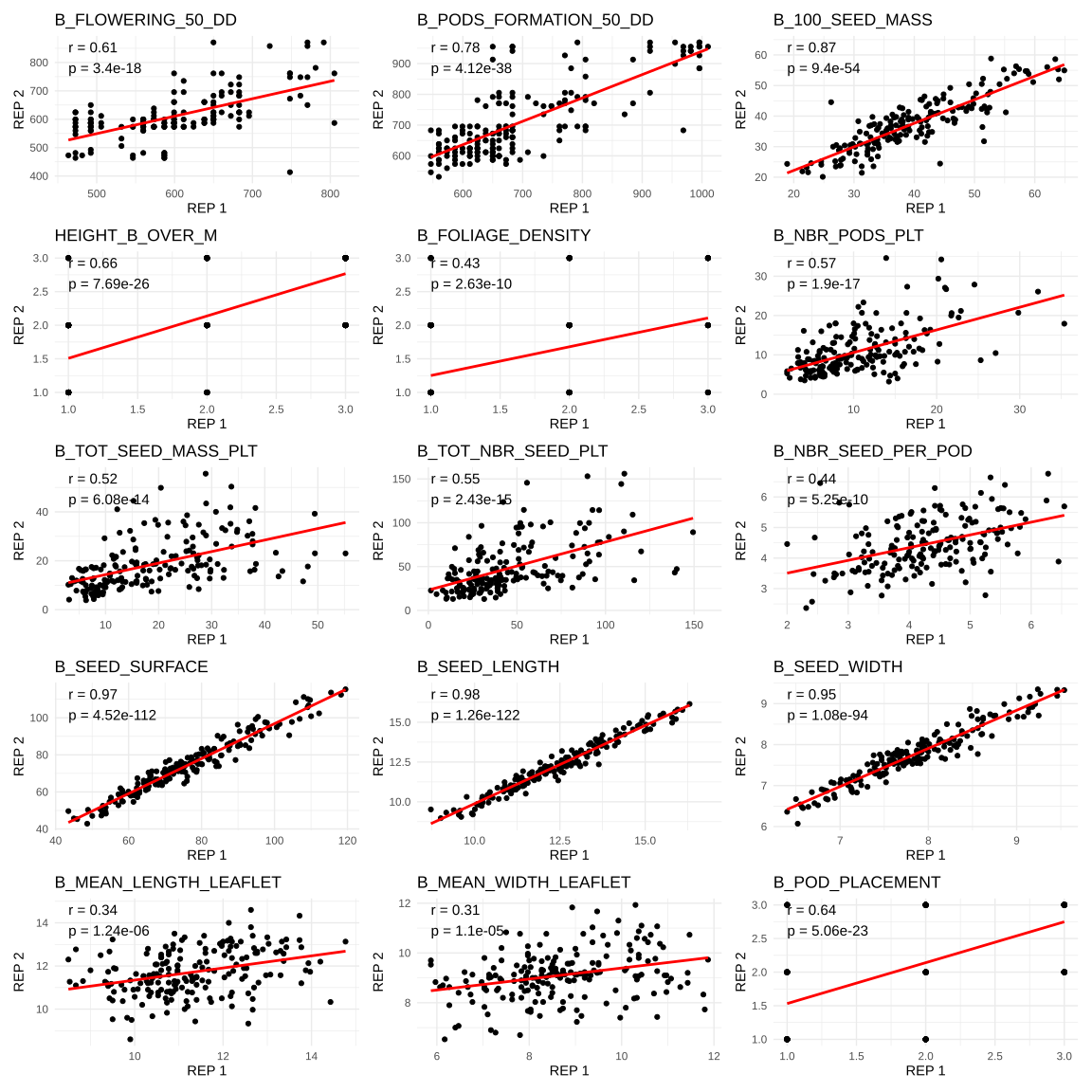

Figure S3. Pearson correlations between block 1 (REP 1) and block 2 (REP 2) of raw data for 15 bean traits (as defined in Table S3) for plants grown in association with the Gandino maize at the Romanian field site in 2022. Red lines represent the linear regression of block 2 on block 1. For each trait, the correlation coefficient (r) and *P*-value (p) are displayed at the top-left of the plot.

**ALT TEXT:** Fifteen scatterplots compare block 1 and block 2 measurements for bean traits in Romania in 2022 with Gandino maize. Most traits show positive relationships between blocks, with especially tight agreement for seed dimensions and more variable agreement for phenology, foliage, and yield traits; each panel reports Pearson r and P.

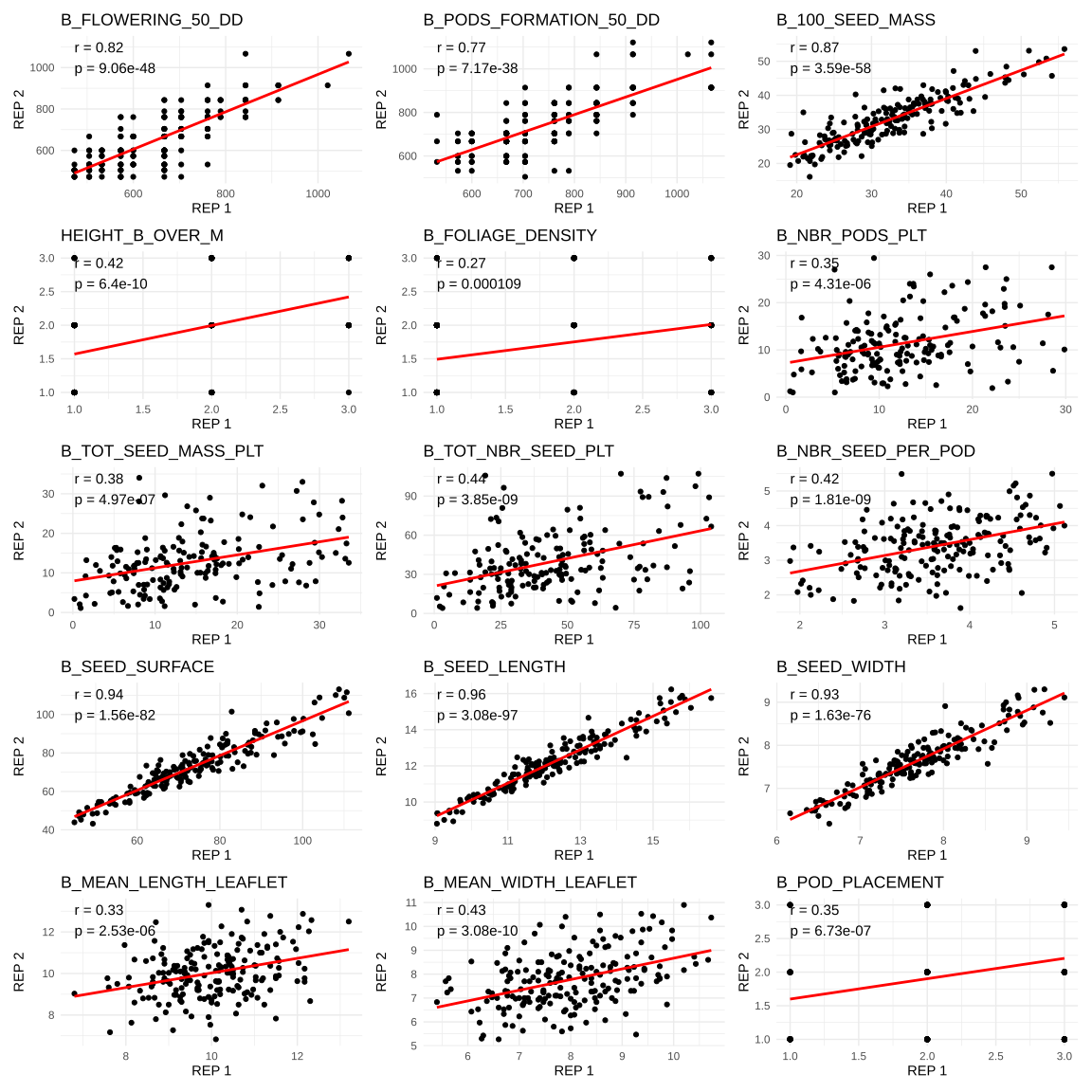

Figure S4. Pearson correlations between block 1 (REP 1) and block 2 (REP 2) of raw data for 15 bean traits (as defined in Table S3) for plants grown in association with the Gandino maize at the Romanian field site in 2023. Red lines represent the linear regression of block 2 on block 1. For each trait, the correlation coefficient (r) and *P*-value (p) are displayed at the top-left of the plot.

**ALT TEXT:** Fifteen scatterplots compare block 1 and block 2 measurements for bean traits in Romania in 2023 with Gandino maize. Positive block-to-block relationships are strongest for seed morphology traits, while several developmental and yield traits show greater scatter; each panel reports Pearson r and P.

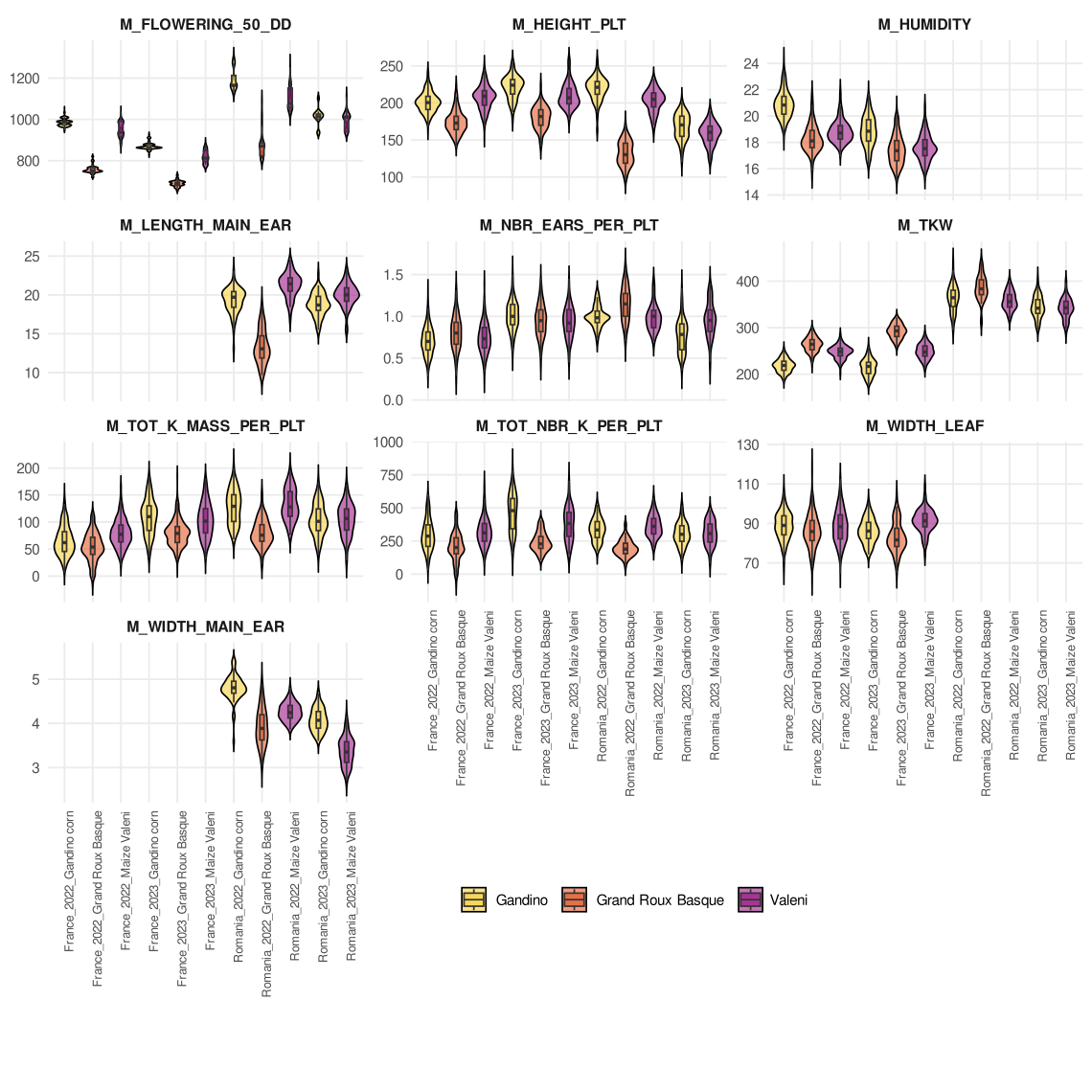

Figure S5. Distributions of maize traits across all experimental fields and for each maize landrace. Distributions are represented as boxplots and violin plots, and the three maize landraces are displayed in different colors. A description of traits is available in Table S3.

**ALT TEXT:** Multi-panel violin and box plots show distributions of maize phenology, architecture, yield, and ear traits for Gandino, Grand Roux Basque, and Văleni across the field experiments. The three landraces differ in the location and spread of several trait distributions.

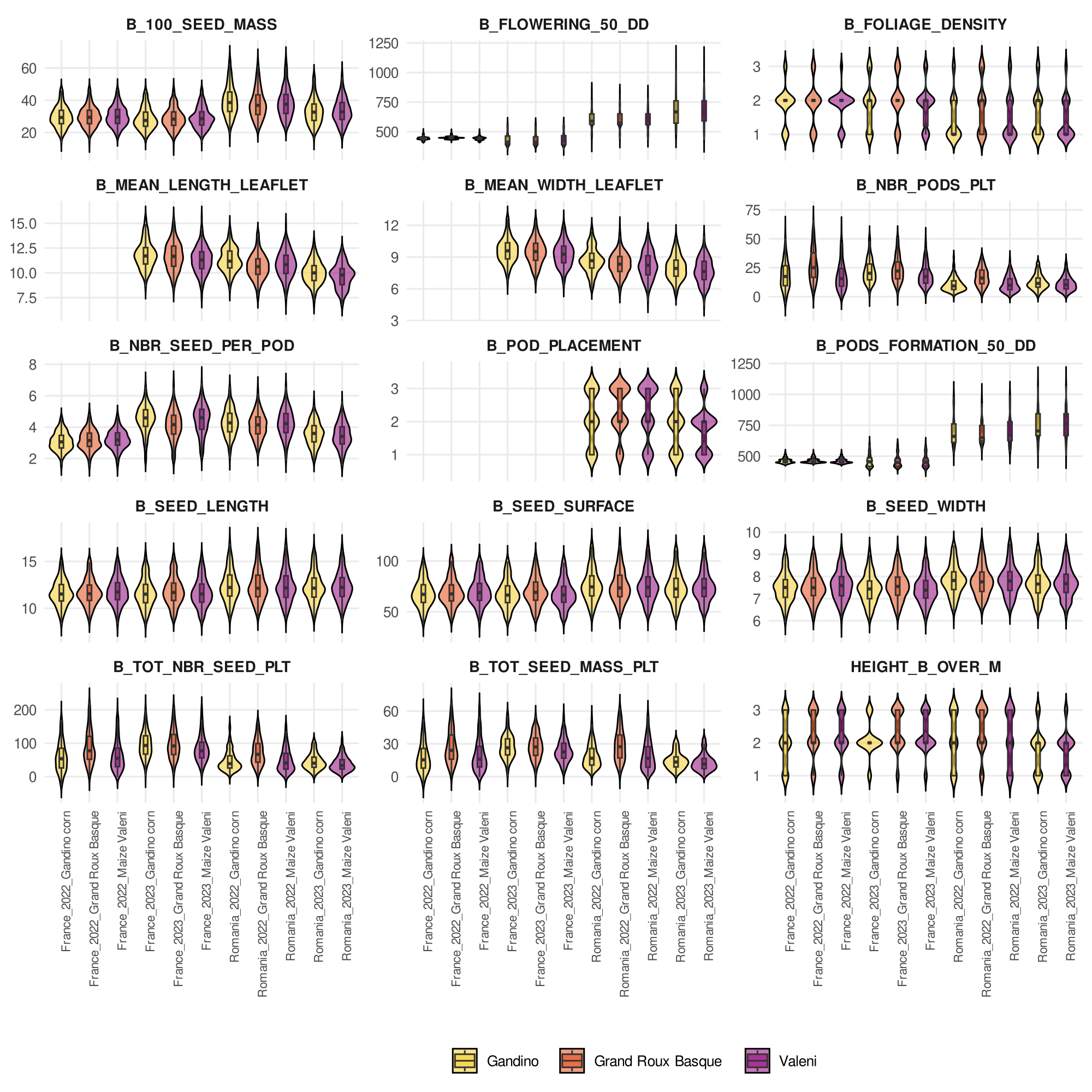

Figure S6. Distributions of bean traits across all experimental fields and for each maize landrace environment. Distributions are represented as boxplots and violin plots, and the three maize landraces are displayed in different colors. A description of traits is available in Table S3.

**ALT TEXT:** Multi-panel violin and box plots show distributions of bean phenology, architecture, yield, and seed traits when grown with Gandino, Grand Roux Basque, and Văleni maize. Several traits shift among maize landrace environments, whereas others show broadly overlapping distributions.

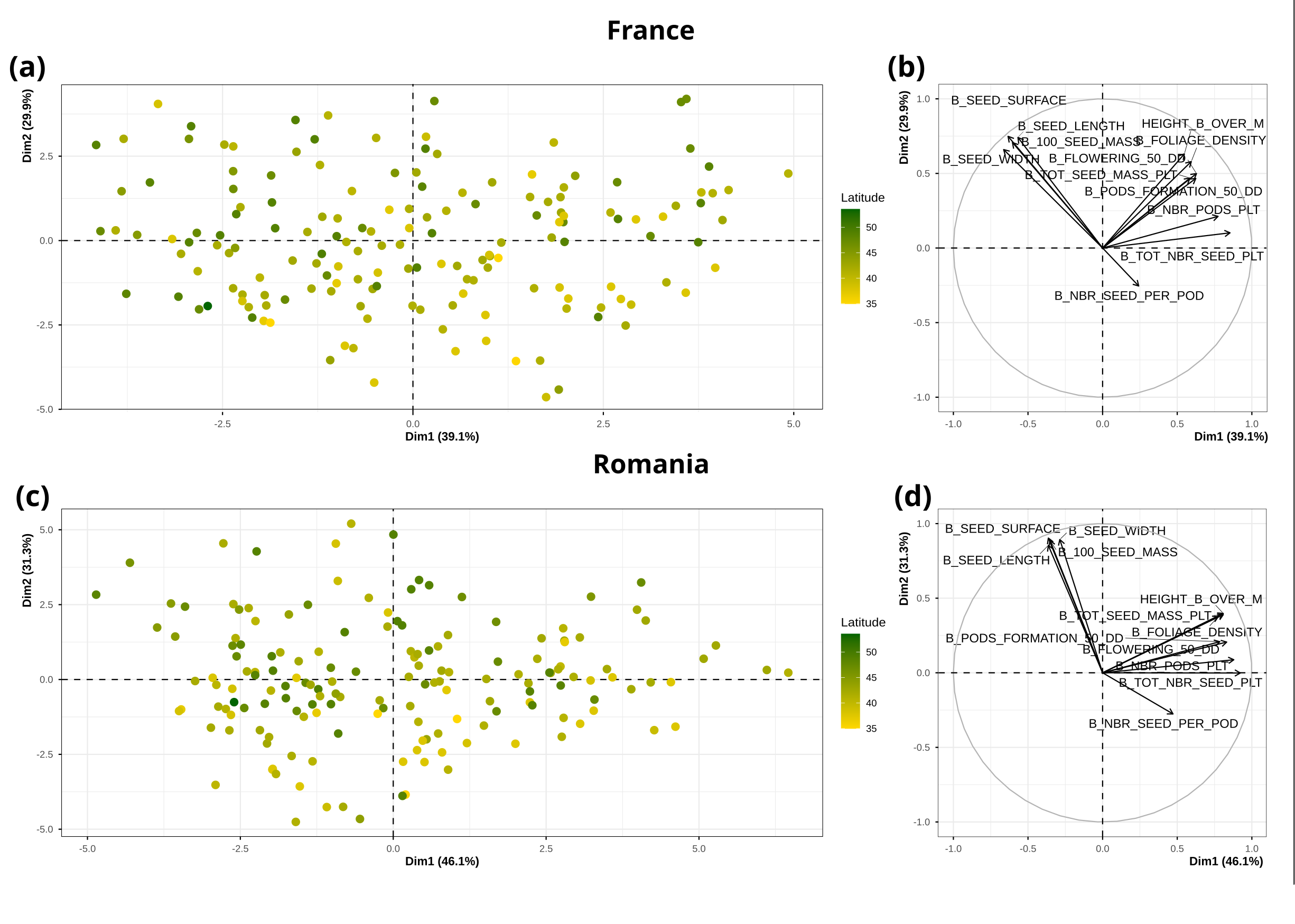
Figure S7. Latitudinal gradients of phenotypic traits measured in France and Romania. PCA was performed on 200 bean lines associated with maize Gandino in the field, using phenotypic data from France (a and b) and from Romania (c and d) corrected for block and year effects (following model M4b). Each PCA includes an individual plot (a and c) and a variable plot (b and d). Colors indicate the latitude of the site of origin for the bean line. A description of traits is available in Table S3.

**ALT TEXT:** Four-panel principal component analysis summarizes bean phenotypes from France and Romania. Individual scores are colored by latitude of origin and accompanied by variable loading plots; latitude forms a gradual color gradient rather than sharply separated clusters.

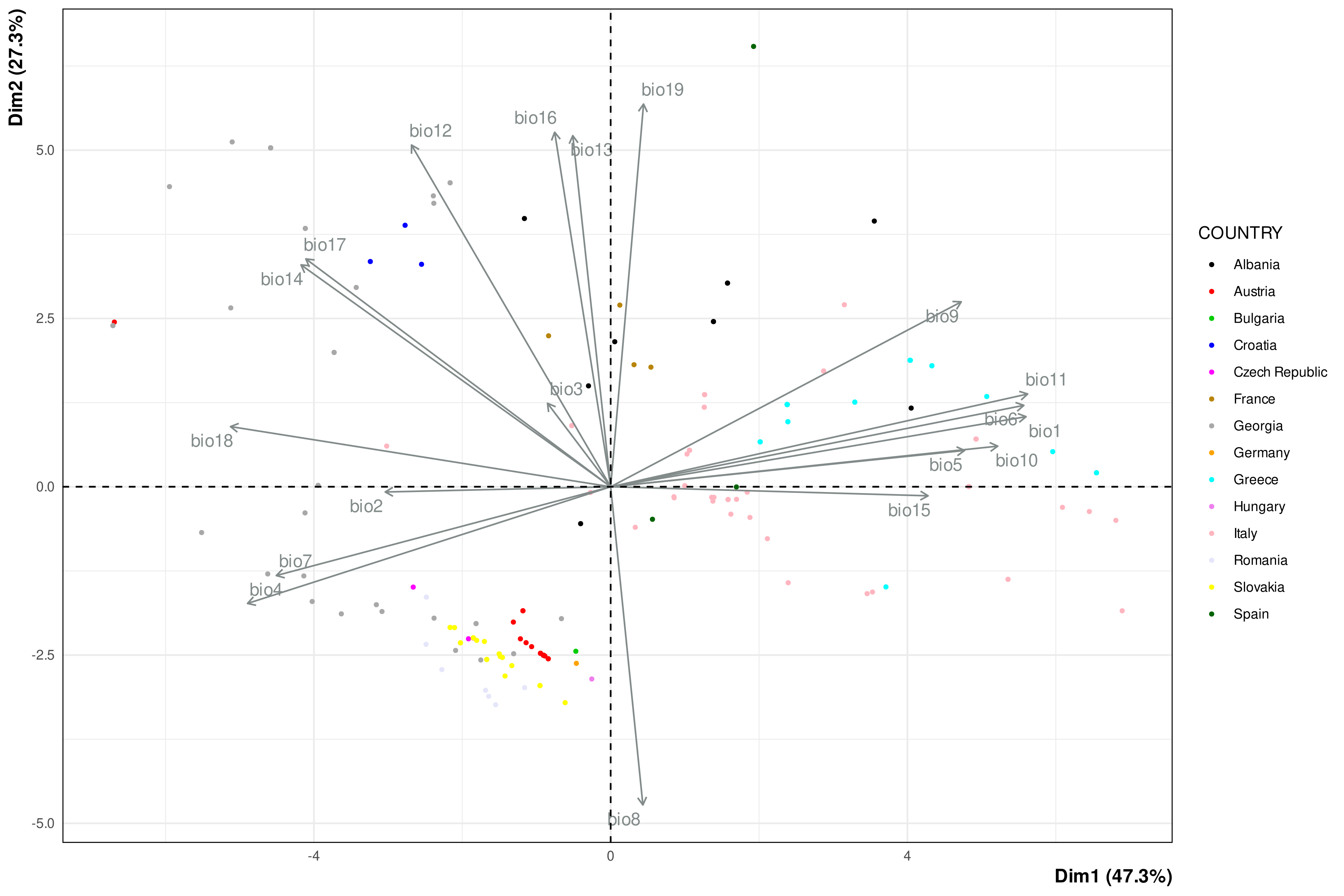

Figure S8. PCA of 183 European bean lines based on 19 bioclimatic variables from the Chelsa database. The figure shows the first two principal components, with a total percentage of variance explained by this plan of 74.6%. Colors correspond to the country of origin of each bean line. The bioclimatic variables are as follows. bio1: mean annual air temperature (°C), bio2: mean diurnal air temperature range (°C), bio3: isothermality (°C), bio4: temperature seasonality (°C/100), bio5: mean daily maximum air temperature of the warmest month (°C), bio6: mean daily minimum air temperature of the coldest month (°C), bio7: annual range of air temperature (°C), bio8: mean daily mean air temperatures of the wettest quarter (°C) bio9: mean daily mean air temperatures of the driest quarter (°C), bio10: mean daily mean air temperatures of the warmest quarter (°C), bio11: mean daily mean air temperatures of the coldest quarter (°C) bio12: annual precipitation amount (kg m^-2^ year^-1^), bio13: precipitation amount of the wettest month (kg m^-2^ month^-1^), bio14: precipitation amount of the driest month (kg m^-2^ month^-1^), bio15: precipitation seasonality (kg m^-2^) bio16: mean monthly precipitation amount of the wettest quarter (kg m^-2^ month^-1^), bio17: mean monthly precipitation amount of the driest quarter (kg m^-2^ month^-1^), bio18: mean monthly precipitation amount of the warmest quarter (kg m^-2^ month^-1^), bio19: mean monthly precipitation amount of the coldest quarter (kg m^-2^ month^-1^).

**ALT TEXT:** Principal component analysis biplot of 183 European bean lines based on 19 bioclimatic variables, with points colored by country of origin and arrows showing variable loadings. The first two components explain 74.6% of variation and reveal partial geographic clustering with substantial overlap among countries.

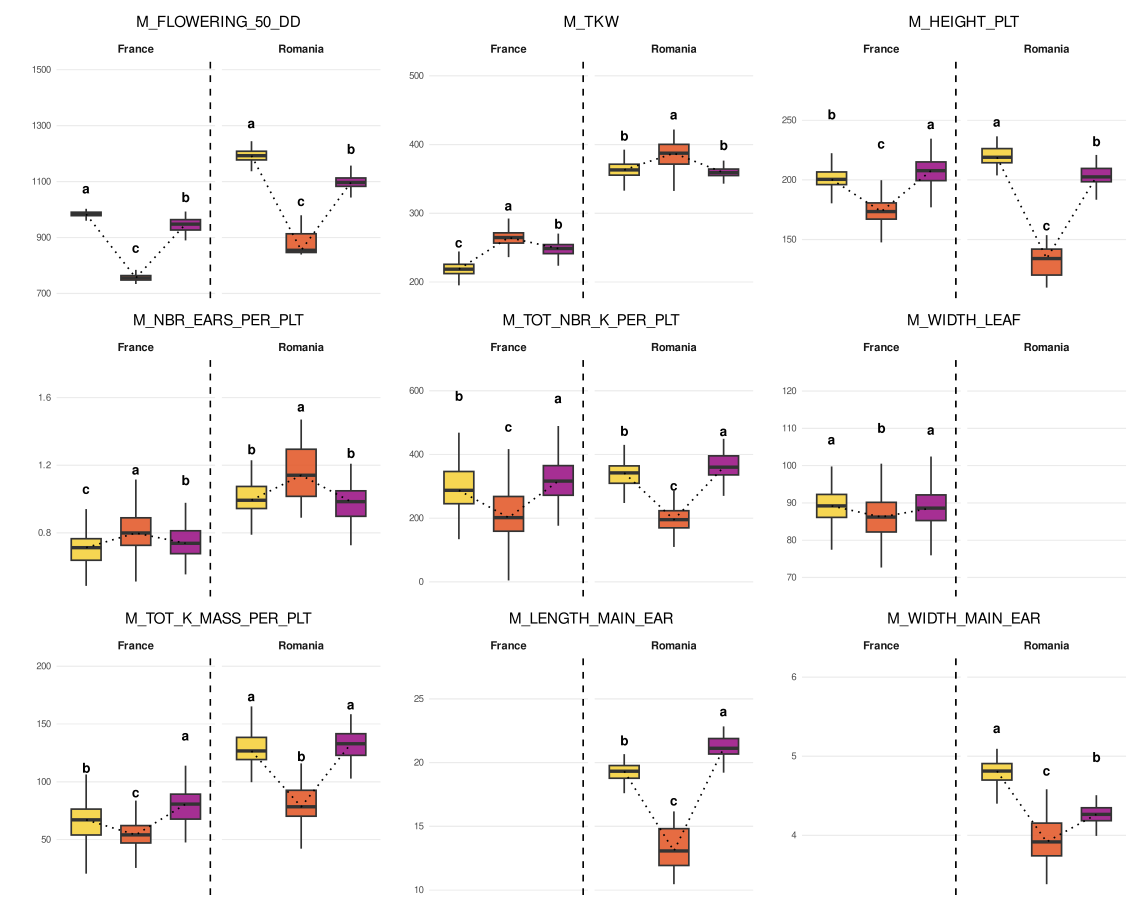
Figure S9. Boxplots of nine maize traits measured in France and/or Romania for the three maize landraces. Figures are based on mean values corrected for the year effects (model M4a). Letters indicate significant differences among maize landraces per country. Traits represented include thermal time to flowering (M_FLOWERING_50_DD), thousand kernel mass (M_TKW), height of plant (M_HEIGHT_PLT), number of ears per plant (M_NBR_EARS_PER_PLT), total number of kernel per plant (M_TOT_NBR_K_PER_PLT), width of the flag leaf (M_WIDTH_LEAF), total kernel mass per plant (M_TOT_K_MASS_PER_PLT), length of the main ear (M_LENGTH_MAIN_EAR) and width of the main ear (M_WIDTH_MAIN_EAR). The dotted lines connect the medians of the three boxplots.

**ALT TEXT:** Nine boxplot panels compare maize traits among Gandino, Grand Roux Basque, and Văleni in France and/or Romania. Letter groupings and dotted median connections show significant landrace differences for several phenology, size, yield, and ear traits.

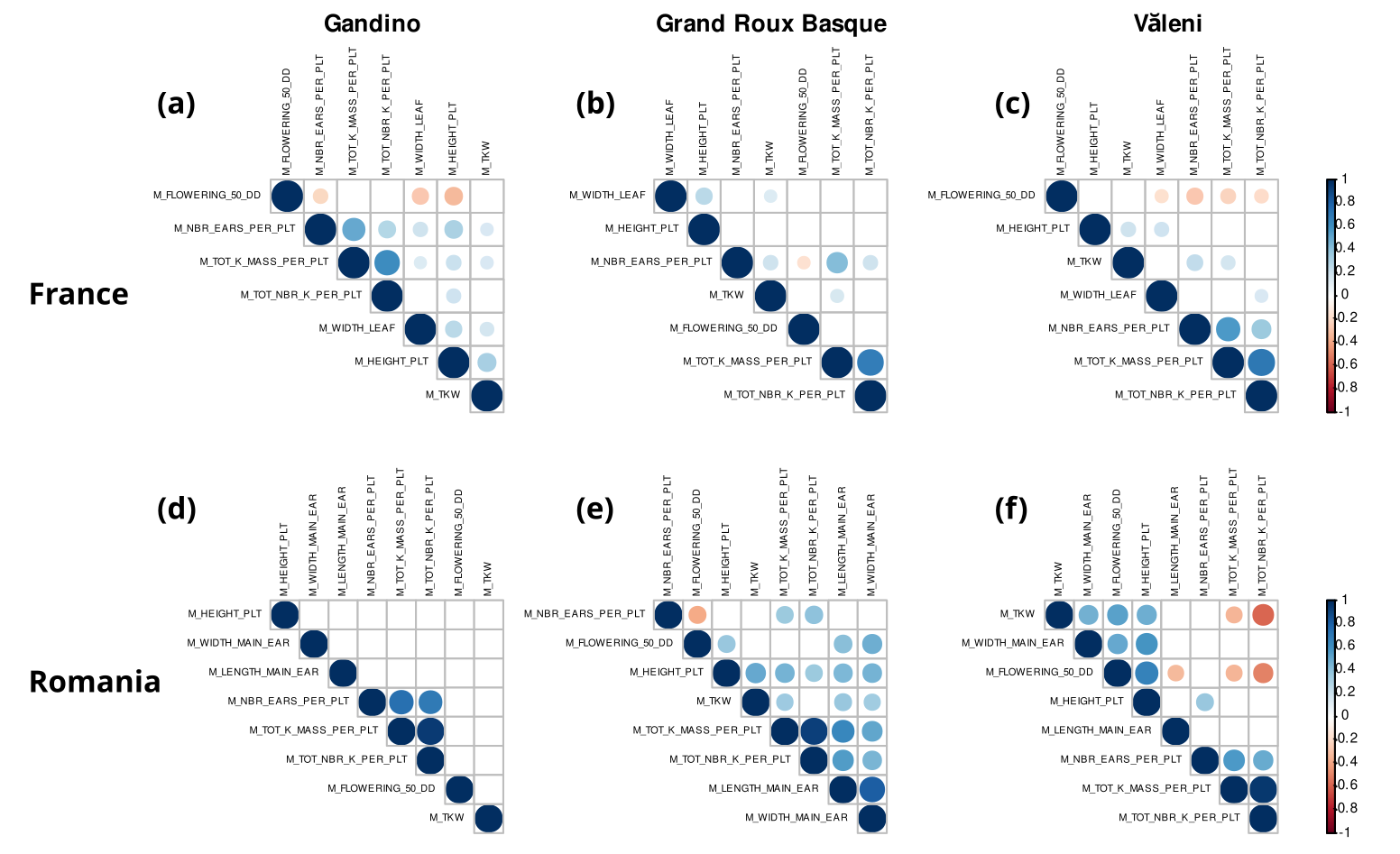

Figure S10. Pearson’s correlation matrices between maize traits for each of the three maize landraces separately (a and d: Gandino; b and e: Grand roux basque; c and f: Văleni) from the French (a-c) and Romanian (d-f) experimental sites. Correlations were calculated on data corrected for the year effects (model M4a). Only significant correlations are reported, based on FDR *q_values*, with a threshold of 5%. A description of traits is available in Table S3.

**ALT TEXT:** Six triangular Pearson correlation matrices show associations among maize traits for each landrace in France and Romania. Significant positive correlations are blue and negative correlations red, with circle size and intensity reflecting correlation strength; correlation patterns differ among landraces and countries.

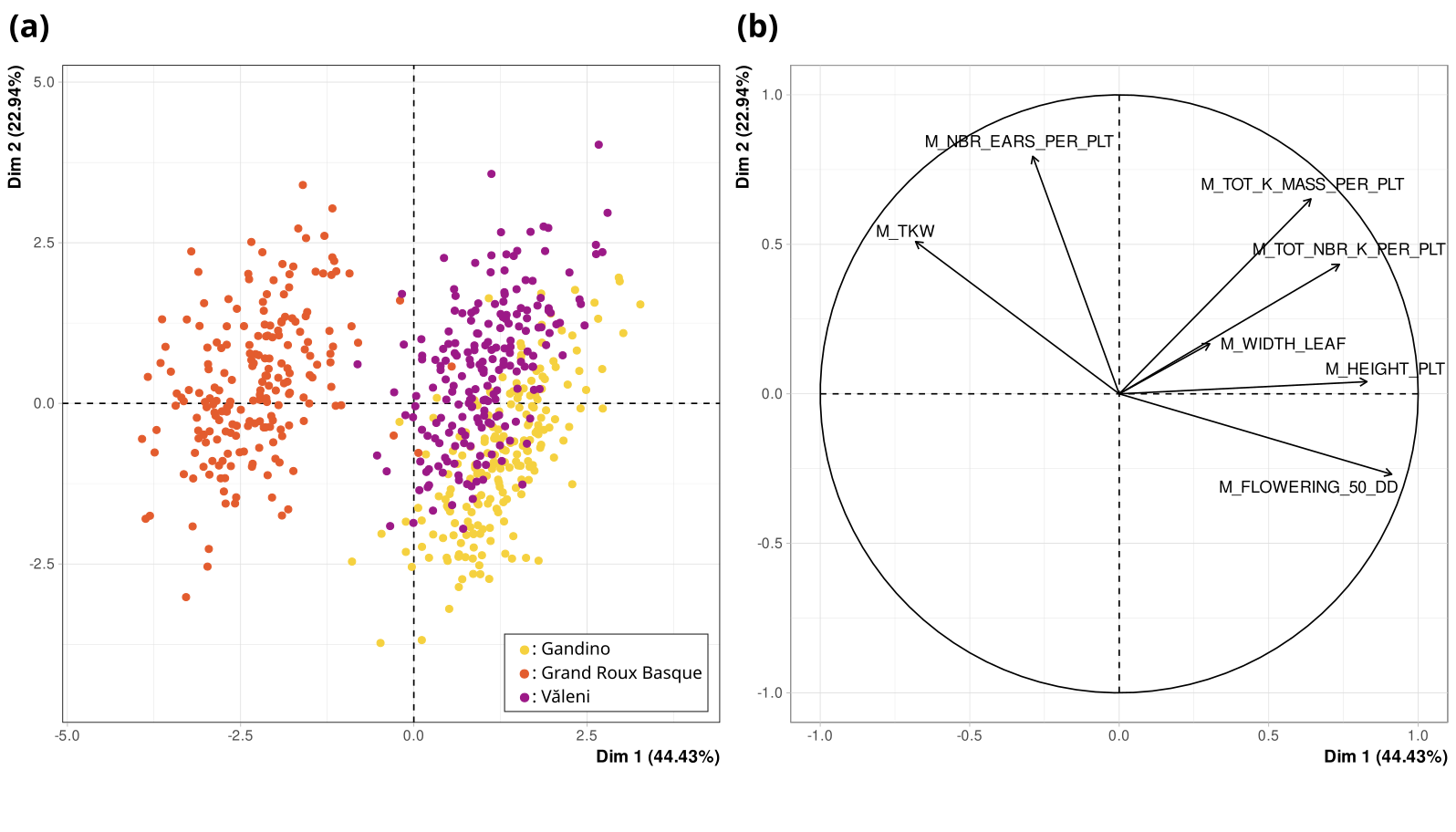
Figure S11. PCA of phenotypic traits measured in France for the three maize landraces : position of individuals (a) and correlation circle of variables (b). The analysis was based on data corrected for block and year effects (model M4a). The plots illustrate the first two principal components, with the percentage of variance explained by each component shown on the corresponding axes. Data are colored following the maize landrace. The seven phenotypic traits include thermal time to flowering (M_FLOWERING_50_DD), height of plant (M_HEIGHT_PLT), width of the flag leaf (M_WIDTH_LEAF), thousand kernel mass (M_TKW), number of ears per plant (M_NBR_EARS_PER_PLT), total number of kernels per plant (M_TOT_NBR_K_PER_PLT), and total kernel mass per plant (M_TOT_K_MASS_PER_PLT).

**ALT TEXT:** Two-panel principal component analysis of maize phenotypes in France. The score plot shows Gandino, Grand Roux Basque, and Văleni forming partially separated clusters, while the correlation circle displays loadings of seven phenology, architecture, and yield traits on the first two components.

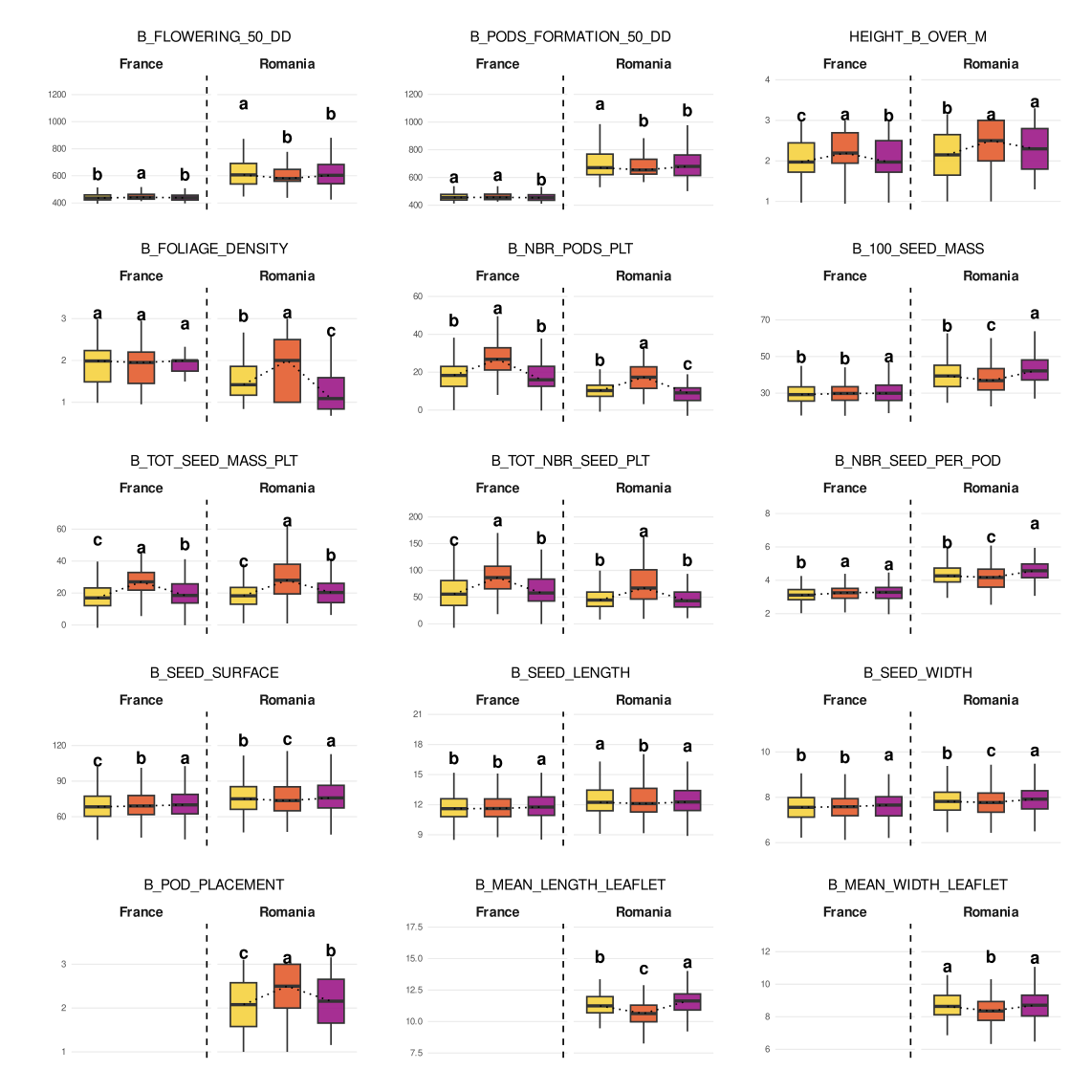

Figure S12. Boxplots of fifteen bean traits measured in France and/or Romania in association with the three maize landraces. Figures are based on mean values corrected for year effects (model M4b). The letters indicate significant differences among maize landraces per country. Traits include thermal time to flowering (B_FLOWERING_50_DD) and to pod formation (B_PODS_FORMATION_50_DD), height of bean over the maize (HEIGHT_B8OVER_M), foliage density (B_FOLIAGE_DENSITY), number of pods per plant (B_NBR_PODS_PLT), 100 seed mass (B_100_SEED_MASS), seed mass per plant (B_TOT_SEED_MASS_PLT), number of seeds per plant and per pod (B_TOT_NBR_SEED_PLT and B_NBR_SEED_PER_POD), seed surface, length and width (B_SEED_SURFACE, B_SEED_LENGTH, B_SEED_WIDTH), pods placement (B_POD_PLACEMENT), leaflet length and width (B_MEAN_LENGTH_LEAFLET and B_MEAN_WIDTH_LEAFLET).

**ALT TEXT:** Fifteen boxplot panels compare bean phenology, architecture, yield, seed, and leaflet traits across Gandino, Grand Roux Basque, and Văleni environments in France and/or Romania. Several traits show significant shifts among associated maize landraces, indicated by letter groups.

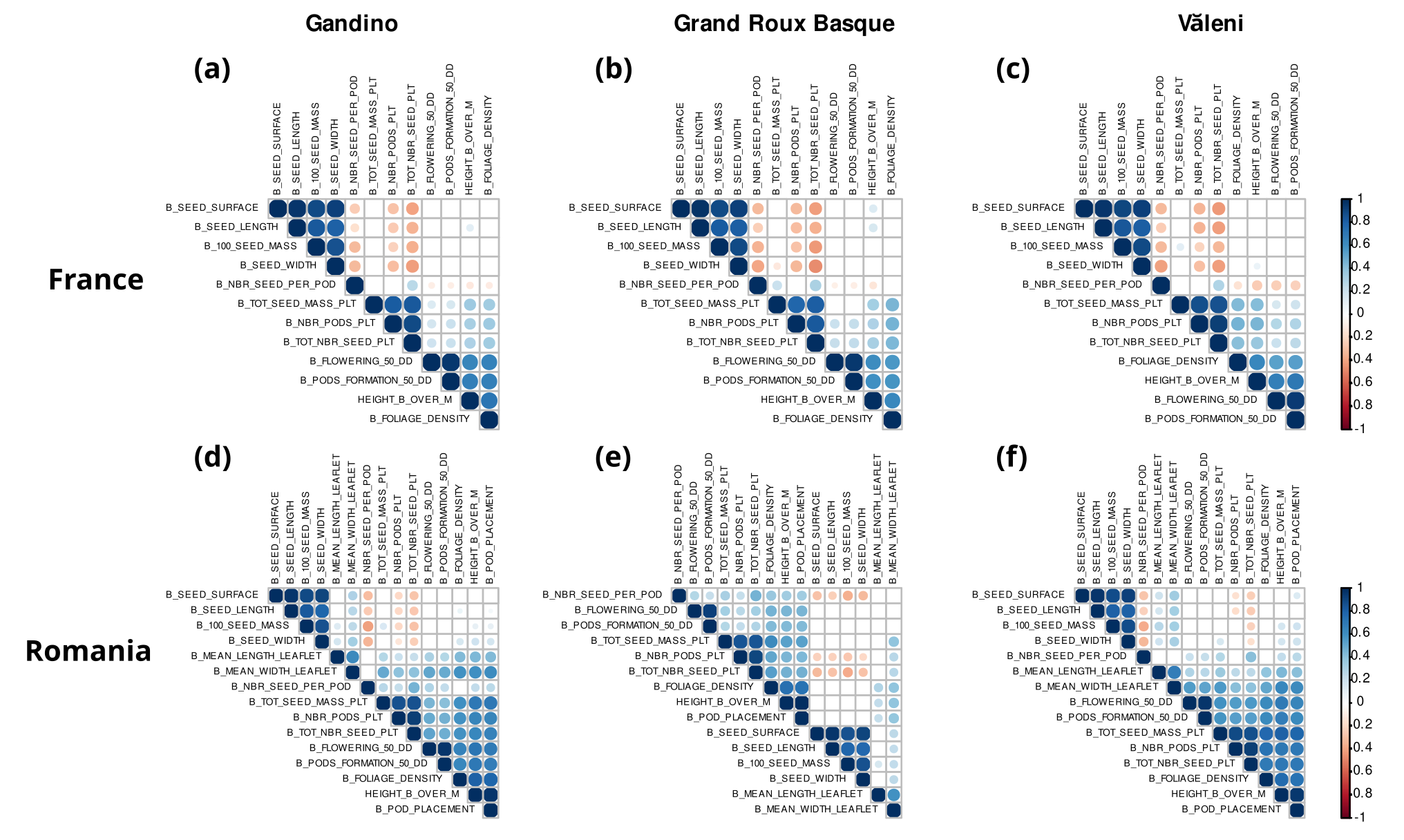
Figure S13. Pearson’s correlation matrices between bean traits for each of the three maize landraces separately (a and d: Gandino; b and e: Grand roux basque; c and f: Văleni) from the French (a-c) and Romanian (d-f) experimental sites. Correlations were calculated on data corrected for the year effects (model M4b). Only significant correlations are reported, based on FDR *q_values*, with a threshold of 5%. A description of traits is available in Table S3.

**ALT TEXT:** Six triangular Pearson correlation matrices summarize relationships among bean traits for each associated maize landrace in France and Romania. Significant positive and negative correlations form broadly similar blocks for some trait groups, with associated maize landrace- and country-specific differences.

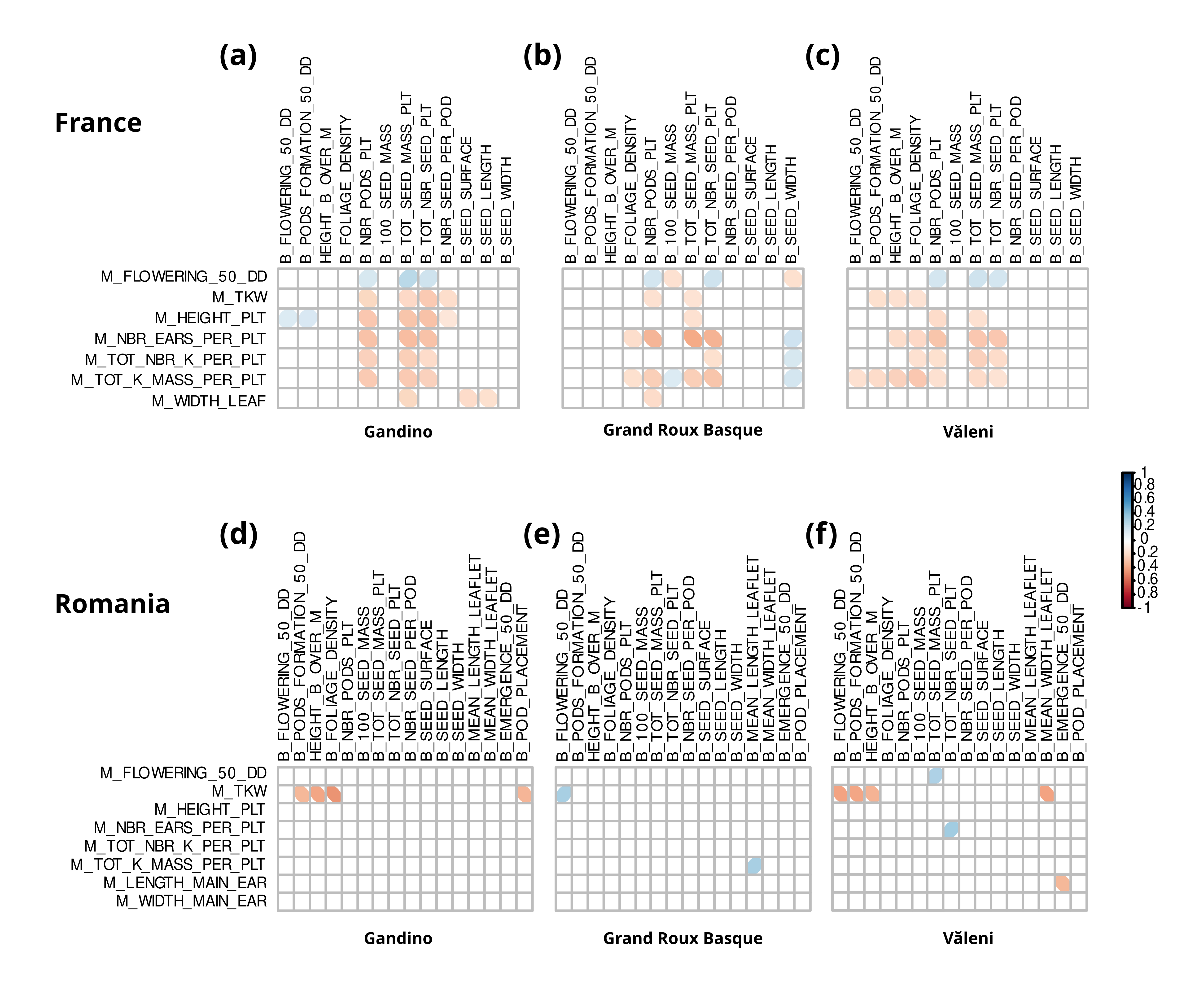
Figure S14. Pearson correlation matrices between maize and bean traits for each of the three maize landraces separately (a and d: Gandino; b and e: Grand Roux Basque; c and f: Văleni) in France (a-c) and in Romania (d-f). Correlations were calculated on data corrected for block and year effects (model M4b). Bean traits are indicated vertically and maize traits horizontally. Positive (blue) or negative (red) correlations are indicated only when FDR q-values were significant at a 5% threshold. A description of traits is available in Table S3.

**ALT TEXT:** Six maize–bean correlation matrices compare each maize landrace in France and Romania. Significant positive and negative correlations between bean traits in rows and maize traits in columns are sparse and vary among associated genotypes and countries.

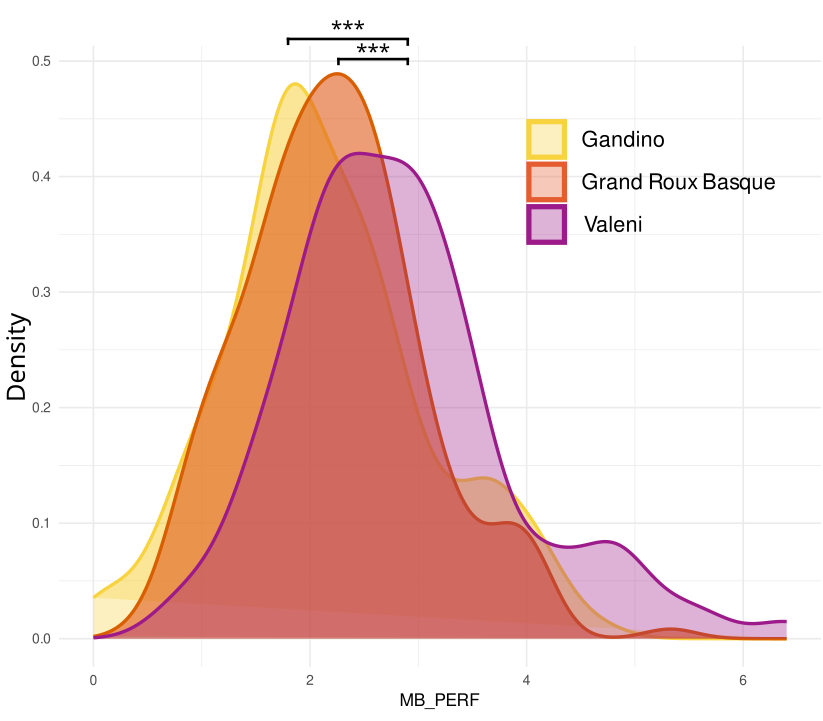

Figure S15. Distribution of the balanced-penalized performance index estimated from the maize kernel and bean seed mass (MB_PERF, Table S3). The data were corrected for the block and the year effects (models M4a and M4b for maize and bean traits, respectively).

**ALT TEXT:** Overlapping density curves show maize-bean corrected balanced-penalized performance for Gandino, Grand Roux Basque, and Văleni. The Văleni distribution is shifted toward higher values relative to Gandino and Grand Roux Basque, with significance brackets indicating differences among distributions.

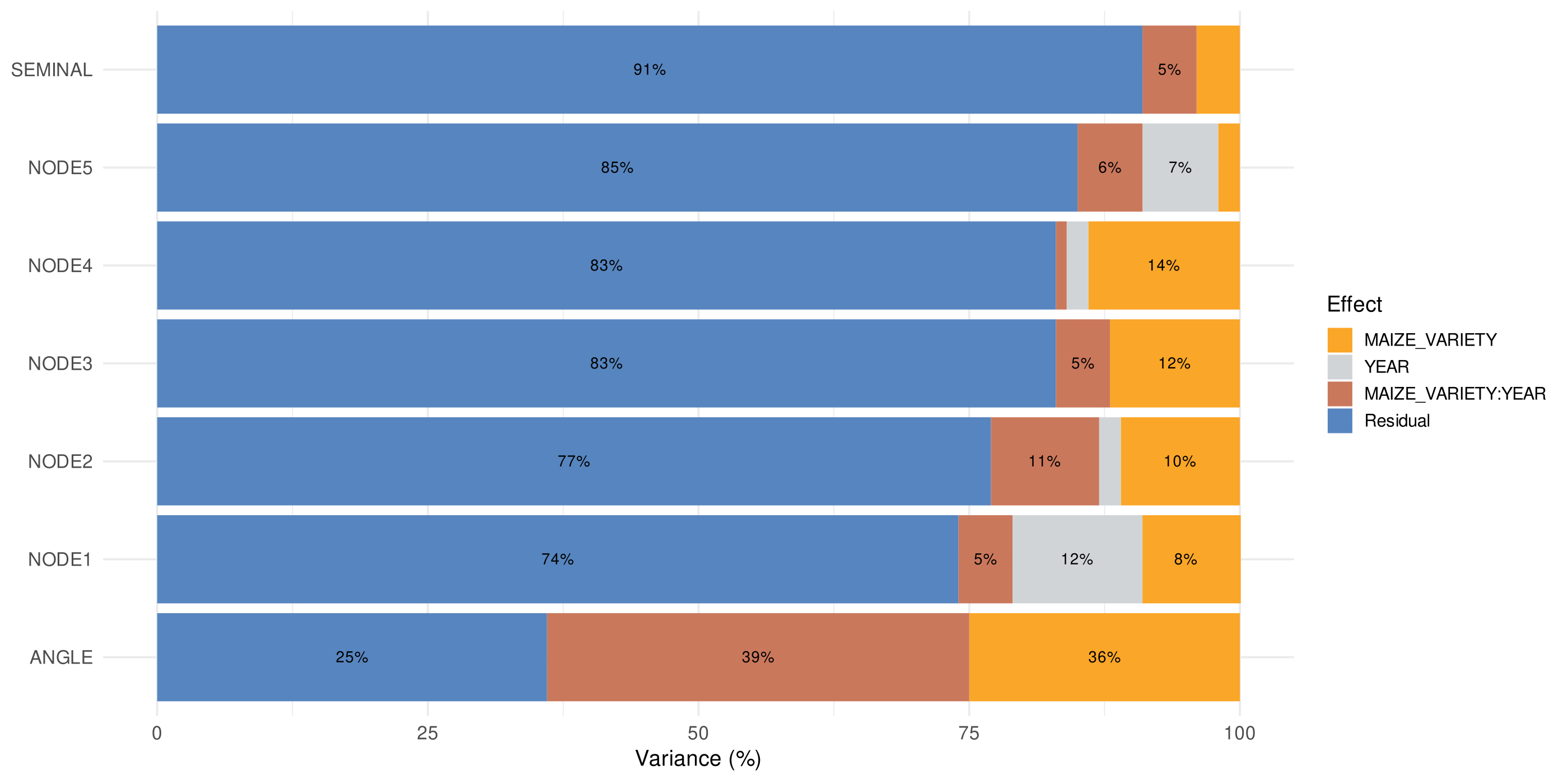
Figure S16. Variance decomposition for the seven maize below-ground traits using model M4a. A description of traits is available in Table S3.

**ALT TEXT:** Horizontal stacked bars partition variance of seven maize below-ground traits among model components. Residual variance is the largest component for most traits, while year, maize landrace and maize-year interaction effects contribute variable smaller fractions, with one trait showing a comparatively large maize and maize-year interaction effects.

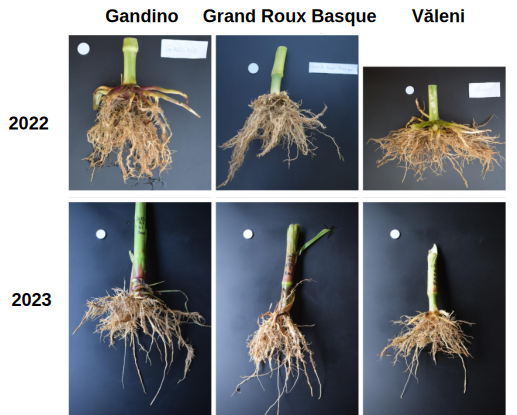

Figure S17. Photos illustrating the three maize landrace root systems in 2022 and 2023.

**ALT TEXT:** Six photographs compare washed root systems of Gandino, Grand Roux Basque, and Văleni maize in 2022 and 2023. Root systems differ visibly among landraces and years in overall spread, branching density, and root mass.

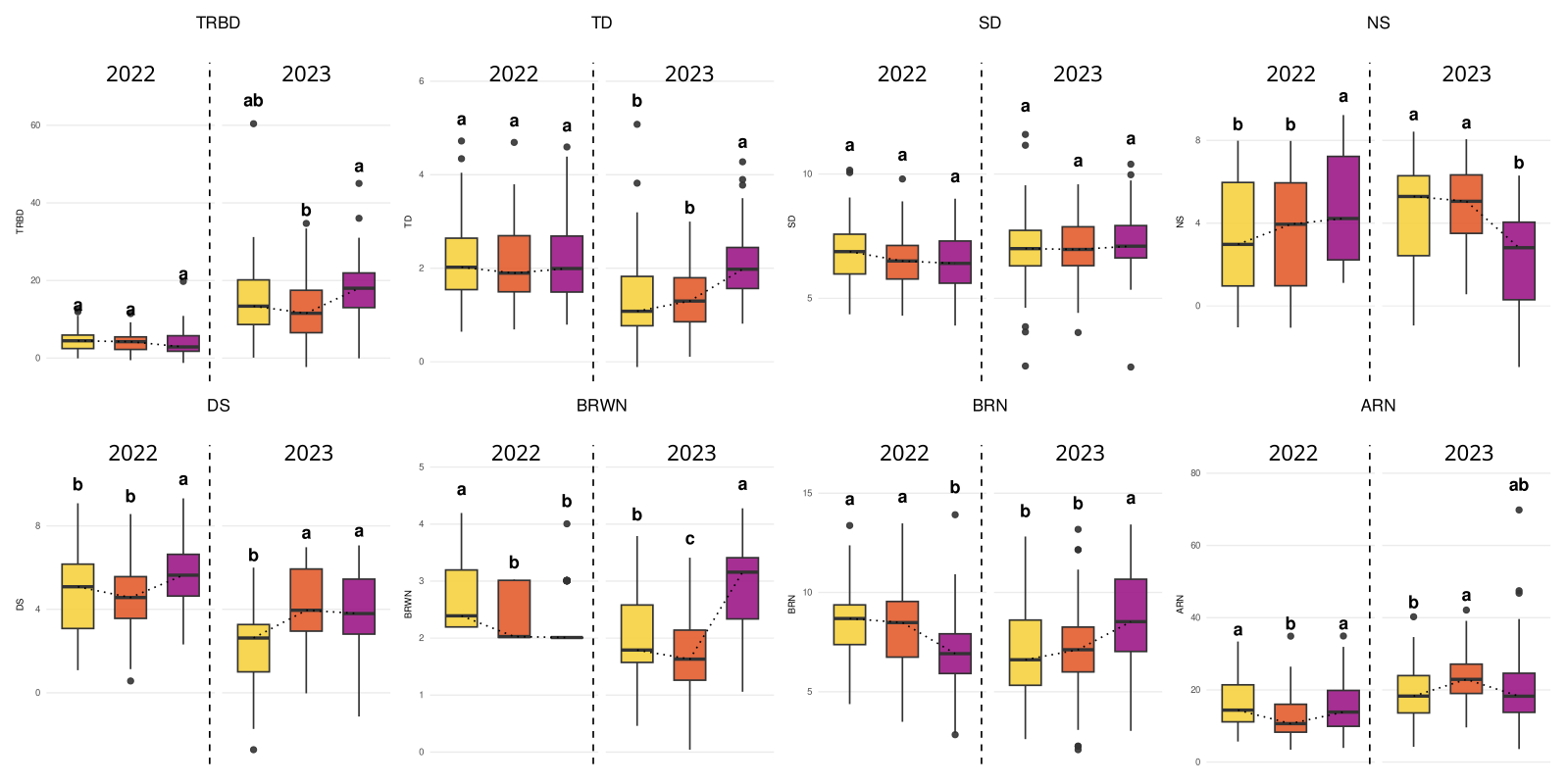

Figure S18. Boxplots of eight manually measured below-ground bean traits in 2022 and 2023 at the French experimental site for the three maize landraces. Figures are based on phenotypic values corrected for the block and manipulator effects (model M2). The letters above the boxplots indicate significant differences among maize landraces for each year separately. A description of traits is available in Table S3.

**ALT TEXT:** Eight boxplot panels compare manually measured bean root traits in 2022 and 2023 across the three maize landraces. Distributions and letter groups show differences among associated maize landraces for several traits, with patterns varying between years.

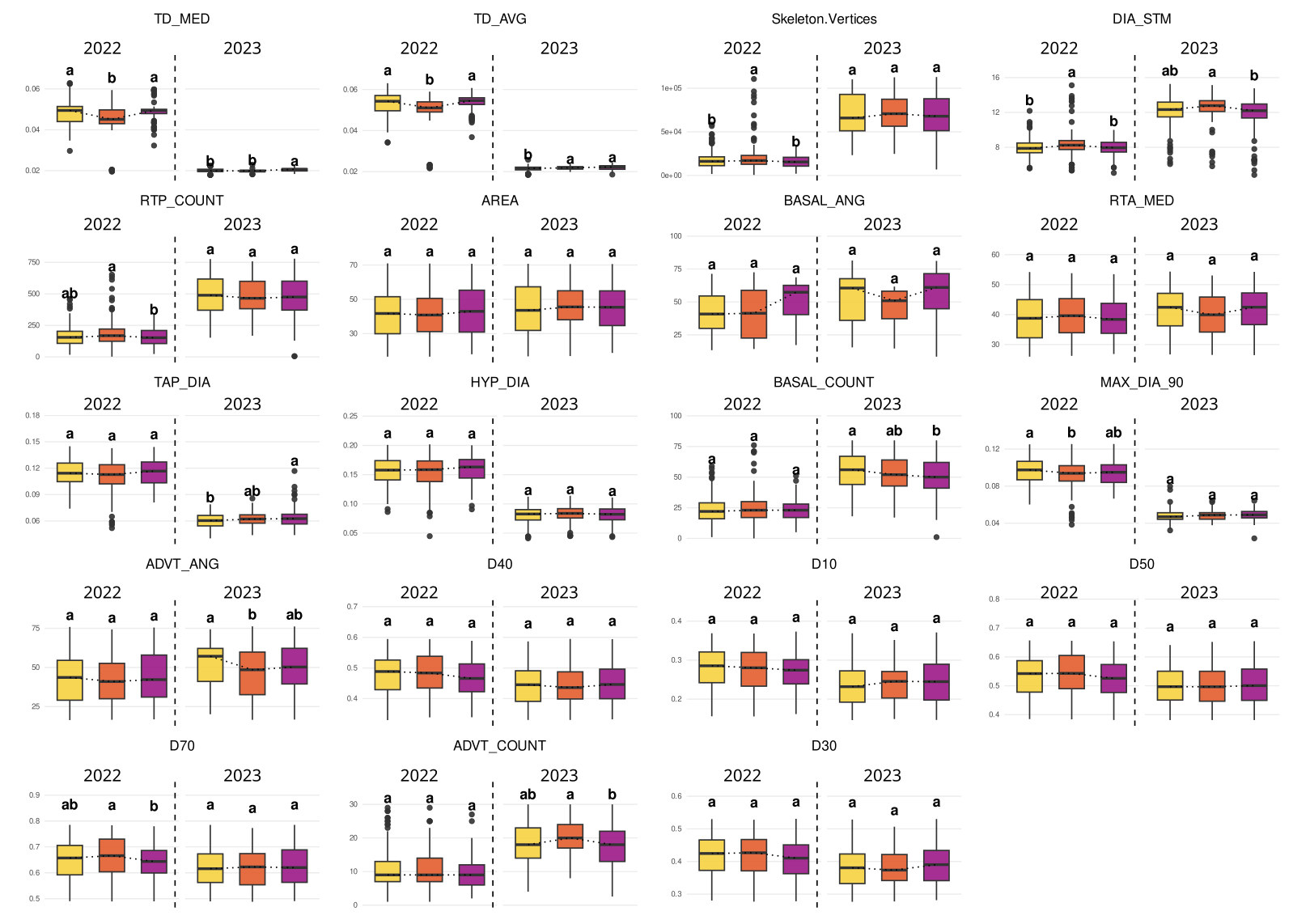

Figure S19. Boxplots of 19 automatically measured below-ground bean traits in 2022 and 2023 at the French experimental site, across the three maize landraces. Values were corrected for block effect (model M2). Letters above boxplots indicate significant differences among maize landraces within each year. A description of traits is available in Table S3.

**ALT TEXT:** Nineteen boxplot panels compare automatic image-derived bean root architecture traits in 2022 and 2023 across Gandino, Grand Roux Basque, and Văleni. Several traits show differences in median and spread among maize environments, with statistical groupings indicated above boxes.
